# Phosphorylation at S345 Converts HIF-2α from a Transcription Factor to an RNA binding protein

**DOI:** 10.64898/2026.04.28.721420

**Authors:** Adam Albanese, Leonard A. Daly, Katrin Prost-Fingerle, Luc G. Elliott, Sally O. Oswald, Gabrielle B. Ecclestone, Gemma Bell, Michael Batie, Daniel J. Rigden, Claire E. Eyers, Joachim Fandrey, Niall S. Kenneth, Violaine See, Sonia Rocha

## Abstract

Oxygen sensing and adaptation are vital for metazoan survival. At the cellular level the response to hypoxia is characterised by a switch in transcriptional programming, primarily mediated by hypoxia-inducible factors (HIFs). HIFs comprise an obligatory heterodimer between an oxygen-sensitive HIF-α subunit and an oxygen-insensitive HIF-1β subunit, which together regulate gene expression in response to hypoxia. Among the three HIF-α proteins in vertebrates (HIF-1α, HIF-2α, and HIF-3α), post-translational modifications (PTMs) play a crucial role in modulating their stability, localisation, and activity. While many modifications have been investigated on HIF-1α, our knowledge of HIF-2α regulation is very limited.

Here, we investigated the function of the recently identified HIF-2α S345 phosphorylation site and demonstrate that this disrupts the interaction between HIF-2α and HIF-1β, thereby silencing the HIF-2 transcriptional response. Additionally, our findings suggest that this modification redirects HIF-2α from its normal transcription factor function towards a role in mRNA fate control, as indicated by mass spectrometry interactome analysis and RNA-immunoprecipitation.

Our discoveries highlight a new regulatory mechanism of HIF-2 activity, where S345 phosphorylation impedes HIF-2α/HIF-1β heterodimer formation, creating a switch in HIF-2α function. As HIF-2α is the only HIF-α isoform with a clinically approved inhibitor, these insights are crucial for advancing therapeutic strategies targeting hypoxia signalling.

## Introduction

Oxygen is a crucial metabolite in numerous cellular processes, including aerobic respiration, where it functions as the terminal electron acceptor in the final stage of the mitochondrial respiratory chain. Consequently, all metazoan life is reliant on molecular oxygen (Kaelin & Ratcliffe, 2008; Taylor & McElwain, 2010), and the ability to perceive, and adjust to changes in oxygen concentration, whether that be at the cellular, tissue, or organismal level, is paramount. Eukaryotes have therefore developed highly sophisticated mechanisms for oxygen sensing and adaptation.

The cellular hypoxia response is primarily controlled by a family of transcription factors called hypoxia-inducible factor (HIF) proteins. The HIF family of proteins is comprised of oxygen-sensitive HIF-α proteins and an oxygen-insensitive HIF-1β (also known as the aryl hydrocarbon receptor nuclear translocator, gene name ARNT) subunit, which form an obligatory heterodimer to form a functional transcription factor (Wang *et al*, 1995). Most vertebrates, including humans, possess three distinct HIF-α proteins (Hara *et al*, 2001). HIF-1α (the most extensively studied HIF-α protein) is expressed across all cell types of vertebrate organisms and is readily degraded in normoxia (Wang & Semenza, 1995). HIF-2α (otherwise termed Endothelial PAS domain protein -1, gene name *EPAS1*) expression displays greater tissue specificity than HIF-1α, being limited to endothelial cells (ECs), cardiomyocytes, kidney fibroblasts, interstitial cells as well as hepatocytes (Tian *et al*, 1997; Zhao *et al*, 2015). Canonically, HIF-2α is involved in the regulation of erythropoietin (EPO) production as well as high-altitude adaptation (Eckardt & Kurtz, 2005; Hanaoka *et al*, 2012; Yi *et al*, 2010). HIF-3α is the least studied HIF-α protein and is postulated to be a negative regulator of the HIF-1α and HIF-2α proteins with multiple splice variants (Gu *et al*, 1998; Makino *et al*, 2001; Makino *et al*, 2002; Maynard *et al*, 2003). HIF-1β is constitutively stable irrespective of oxygen tension, with active roles in both normoxia and hypoxia, but only forms the HIF heterodimeric complex upon stabilisation of HIF-α (Wang *et al*., 1995).

HIF-α stabilisation during hypoxia is regulated by post-translational modifications (PTMs). Prolyl hydroxylation of HIF-α within their respective oxygen-dependent degradation (ODD) domains occurs through the action of a family of prolyl hydroxylase domain-containing proteins (PHDs) (Bruegge *et al*, 2007; Fandrey *et al*, 2006; Masson *et al*, 2001), forming a recognition interface for interaction with the VHL E3 ubiquitin ligase complex, resulting in polyubiquitylation of HIF-α and targeted protein degradation via the 26S proteasome (Bruick & McKnight, 2001; Epstein *et al*, 2001; Ivan *et al*, 2001). However, when cells become hypoxic, oxygen is no longer available to act as a cofactor for the PHDs, reducing HIF-α prolyl hydroxylation, stabilising HIF-α where it can accumulate in the nucleus, heterodimerising with HIF-1β via their Per-Arnt-Sim (PAS) domains. HIF is then capable of binding hypoxia response elements (HRE) of genes to initiate hypoxic gene expression (Wenger *et al*, 2005). HIF hydroxylation is not solely regulated by the PHD proteins, but also by the factor inhibiting HIF (FIH) (Lando *et al*, 2002; Mahon *et al*, 2001). The C-terminal activation domain (CTAD) is regulated in an oxygen-dependent manner via FIH-mediated asparagine hydroxylation which suppresses co-activator CBP/p300 binding. When oxygen tension is further reduced, FIH no longer modifies HIF-α, allowing for CBP/p300 co-activator binding and enhanced target gene induction (Lando *et al*., 2002).

Despite HIF-1α being the most studied HIF-α protein, HIF-2α remains the only HIF transcription factor with a clinically approved direct inhibitor, Belzutifan (Deeks, 2021). While Belzutifan was initially marketed as a novel treatment for von Hippel–Lindau tumour suppressor (VHL) disease, a genetic disorder that predisposes individuals to tumour development, it is now an approved treatment option for renal cell carcinoma (RCC) (Kao *et al*, 2023). Beyond this, the co-administration of HIF-2α inhibitors with other therapies is postulated to improve treatment responsiveness in renal cancers (Choueiri *et al*, 2025).

While the primary mode of HIF-α regulation occurs on the post-translational level, involving the canonical PHD/VHL-axis, additional modes of regulation are well documented at the transcriptional, translational, and post-translational levels (Albanese *et al*, 2020). PTMs increase proteome complexity by altering protein stability, activity, localisation, and binding partners (Lee *et al*, 2023; Wang *et al*, 2022). HIF-1α and HIF-2α are no exceptions. While many PTM sites have been identified on HIF-1α and HIF-2α (reviewed in (Albanese *et al*., 2020)), a significant proportion of these are yet to be functionally characterised and the current knowledge surrounding HIF-2α PTM-dependent regulation is very limited (Daly *et al*, 2021).

This investigation explores the role of the hypoxia-specific HIF-2α S345 phosphorylation site (Daly *et al*., 2021). Phosphomimetic mutation of HIF-2α S345 severely compromises HIF transcriptional activity by selectively abrogating heterodimerisation with HIF-1β. Interestingly, this phosphomimetic mutation also enhances HIF-2α binding to specific RNAs. These data thus point to a potential signalling switch in HIF-2α function via phosphorylation at S345, revealing a regulatory mechanism that governs these important functions of HIF-2α.

## Results

### HIF-2α S345 phosphomutants impair HIF activity in an artificial reporter assay

Phosphorylation of HIF-2α on S345 was initially identified as a hypoxic-specific event in an unbiased mass spectrometry (MS) screen exploring the effects of oxygen tension on HIF-2α (and HIF-1α) interactomes and PTM status (Daly *et al*., 2021).S345 lies immediately C-terminal to the PAS-associated C terminal (PAC) motif, which is thought to contribute to PAS domain folding and by extension, HIF heterodimerisation and transcriptional activity (Figure 1A) (Ponting & Aravind, 1997). However, the functional ramifications of this phosphorylation event are currently unknown. Interestingly, conservation analysis of S345 across >200 vertebrates revealed this site to be invariant, with minimal variation also in nearby residues (Figure 1A), thus suggesting conservation of function.

**Figure 1.**
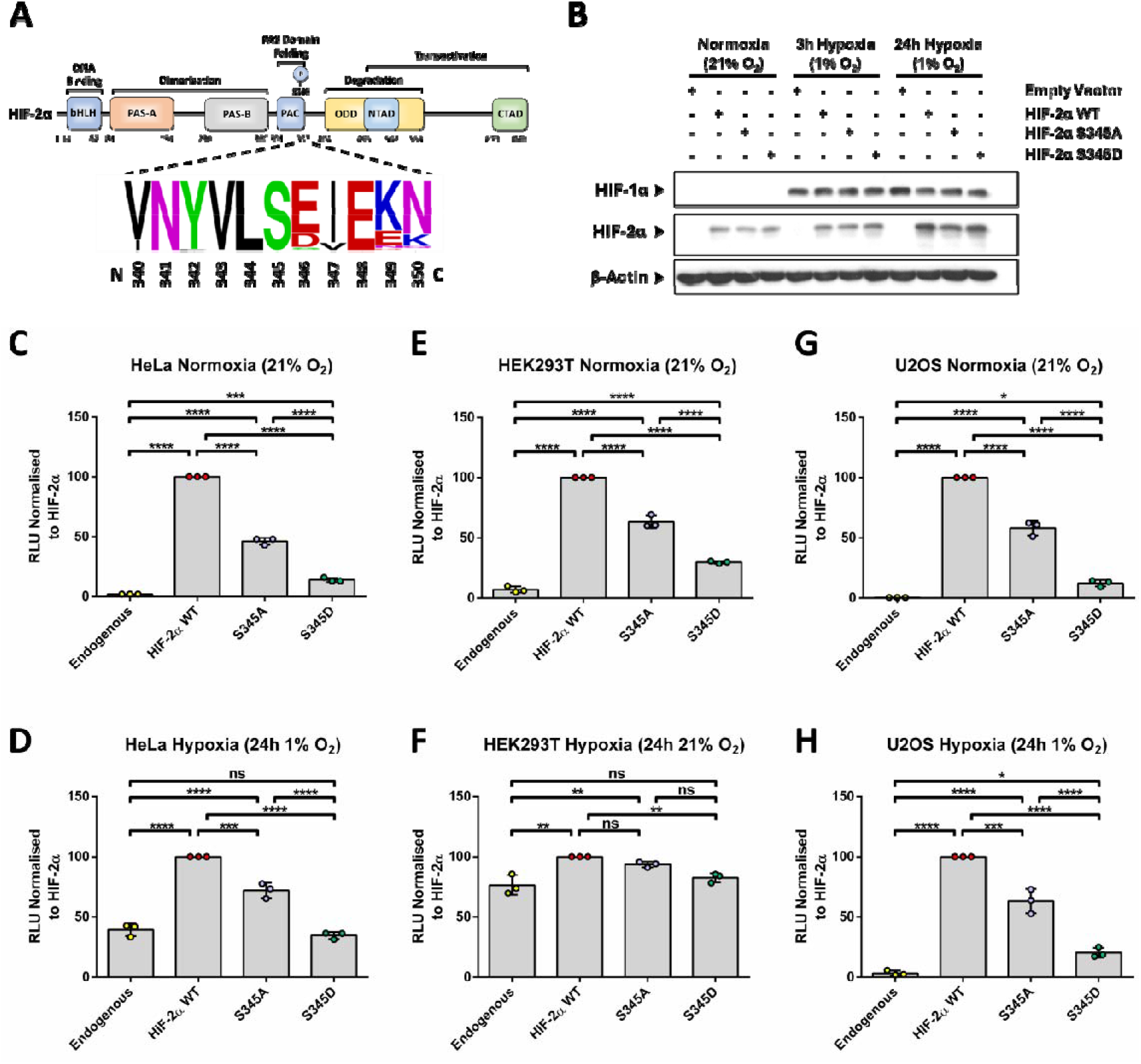
Characterisation of HIF-2α S345 mutants. **(A)** Large-scale MUSCLE protein sequence alignment (>200 vertebrate full-length HIF-2α protein sequences) with schematic focused on the human HIF-2α protein sequence (Q99814) S345 phosphosite ±5 residues. WEBLOGO plot for the frequency of amino acid substitutions at each residue within the S345 consensus sequence (Crooks et al., 2004). (**B**) Representative Western blot for the protein expression levels of HIF-2α S345 phosphomutants. HeLa cells transiently exogenously expressing HA-Clover HIF-2α wild-type (WT), S345A or S345D were exposed to normoxia (21 % O_2_), 3-hour or 24-hour hypoxia (1 % O_2_) (*n* = 3). HeLa cells were transfected for a low expression HIF-2α WT, S345A or S345D construct and HRE-luciferase construct and then exposed to (**C**) normoxia (21 % O_2_) or (**D**) 24-hour hypoxia (1 % O_2_). Comparable luciferase reporter assays were also carried out in (**E** and **F**) HEK293T and (**G** and **H**) U2OS cells exposed to either normoxia or hypoxia (21 % and 1% O_2_, respectively). Luciferase activity was measured using an HRE-luciferase reporter assay. Luminescence levels were normalized to the HIF-2α condition (*n* = 3 independent experiments). ± Standard deviation error bars are reported. One-way ANOVA Tukey post-hoc tests were performed to determine significance (*p < 0.05; ns, not significant).

To investigate the functional consequences of HIF-2α S345 phosphorylation, phosphonull (S345A), phosphomimetic (S345D) and wild-type HIF-2α constructs were transiently transfected in HeLa cells. When exogenously expressed, HIF-2α wild-type, S345A and S345D retained comparable protein expression levels in HeLa cells during normoxia (21% O_2_) (Figure 1B), with only a slight inconsistent reduction in the levels of S345A. Moreover, protein expression levels of the HIF-2α wild-type/S345A/S345D increased following either 3- or 24-hours hypoxia (1% O_2_), with similar levels of expression between each of the HIF-2α variants. To assess whether HIF-2α S345 phosphorylation could impact HIF transcriptional activity, a HIF-dependent luciferase reporter assay was performed in HeLa cells (Figures 1C-D). Co-expression of HIF-2α wild-type along with the HRE reporter demonstrated significant increases in reporter activity beyond the endogenous levels at 21% O_2_ which increased further following 24-hours hypoxia. The activity of both S345A and S345D was significantly reduced compared to wild-type protein in both hypoxia and normoxia, with the activity of the S345D variant being markedly low under both oxygen tensions. Notably, the activity of S345D HIF-2α was not increased above endogenous levels following 24-hours hypoxia. Similar findings were observed in two additional cell lines, U2OS and HEK293T, with S345D HIF-2α having almost no transcriptional activity as determined by the reporter assays (Figures 1E-H). Together this suggests that the S345D phosphomimetic has severely reduced HIF transcriptional activity but does not act as a dominant negative mutant.

### HIF-2α S345D abolishes endogenous HIF target gene induction

While informative, the luciferase assays do not contain the correct chromatin environment to evaluate endogenous transcriptional function. It was therefore important to determine if the expressed HIF-2α mutants altered endogenous target gene expression. In HeLa cells, by Western blot, the expression levels of both PHD3 and NDRG1 increased upon overexpression of either HIF-2α wild-type or S345A, relative to the endogenous control (Figure 2A). However, this was not the case for HIF-2α S345D whose expression was more comparable to that of the endogenous control. This increase in target protein expression was mirrored at the mRNA expression level (as determined by RT-qPCR), with significant increases being observed for NDRG1, PHD3 and GLUT3 transcripts following exogenous expression of either HIF-2α wild-type or S345A compared to endogenous control (Figure 2B-G). However, for HIF-2α S345D no significant change was observed in PHD3 and GLUT3 transcript abundance beyond control levels, irrespective of normoxia or 24-hours hypoxia. Similar data was also obtained in HEK293T cells (Sup. Figure 1). Together this suggests that despite consistent expression levels, HIF-2α S345D is unable to induce transcription.

**Figure 2.**
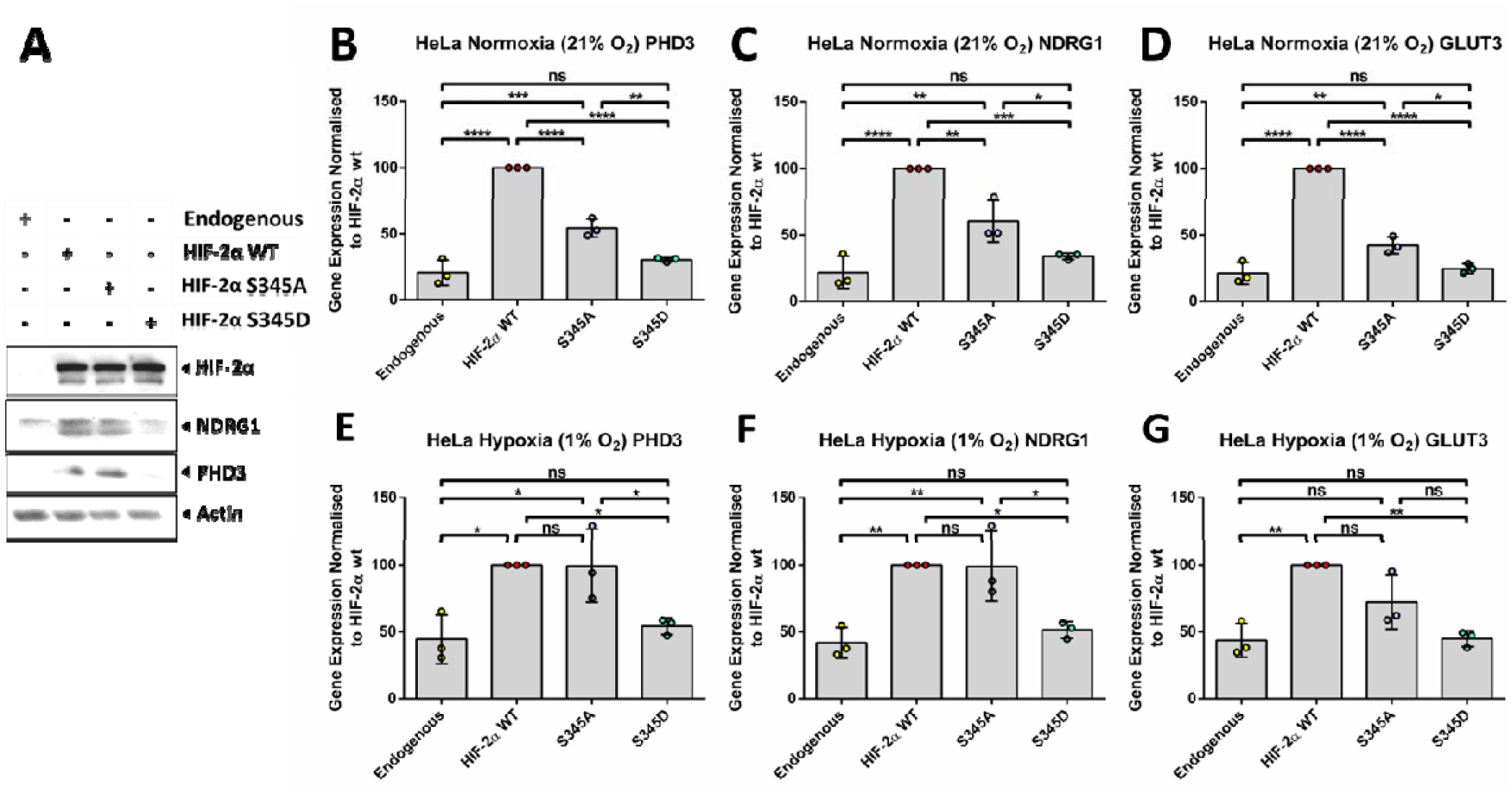
HIF-2α S345D reduces HIF induction of endogenous target genes at protein and gene level. (**A**) Representative Western blot of HeLa cells transiently transfected with either empty vector, HA-Clover HIF-2α, HA-Clover HIF-2α S345A or HA-Clover HIF-2α S345D for high level overexpression. Protein expression levels were determined for HIF-2α, HIF targets NDRG1 and PHD3, as well as β-actin as a loading control. (**B, C, D, E, F** and **G**) HIF target induction in HeLa cells were likewise determined at the gene level via RT-qPCR for (**G** and **H**) PHD3, (**I** and **J**) NDRG1 and (**K** and **L**) GLUT3, in normoxia and hypoxia (21% and 1% O_2_, respectively). All experiments were carried out as *n* = 3 independent experiments. Gene expression levels were normalised to HIF-2α WT. ±Standard deviation error bars are reported. One-way Anova Tukey post-hoc test was performed to determine significance (*P < 0.05; ns, not significant).

### Mimicking HIF-2α S345 phosphorylation impairs interaction with HIF-1β

Site-specific phosphorylation of HIF-α, like other transcription factors (e.g. RelA), can control its nucleocytoplasmic shuttling and transcriptional activity (Gkotinakou *et al*, 2019; Lanucara *et al*, 2016; Mylonis *et al*, 2008; Mylonis *et al*, 2006). To identify the mechanism behind the HIF-2α S345D-induced activity change, sub-cellular protein localisation and expression levels were assessed by fixed cell fluorescent microscopy (Sup. Figure 2A). In line with protein levels demonstrated by Western blot, protein accumulation between wild-type and S345D remained consistent with no significant changes in fluorescent intensities and nuclear/cytoplasmic localisation (Figure 1B and Sup. Figure 2B).

For HIF to act as a transcription factor, HIF-α must bind to HIF-1β (Wang *et al*., 1995). HIF dimer formation occurs through multiple interfaces in the PAS-A, PAS-B, and bHLH domains (Figure 1A). Moreover, casein kinase 1δ–dependent phosphorylation of HIF-1α S247, residing within the PAS-B domain, is reported to impair HIF complex formation, decreasing transcriptional activity (Kalousi *et al*, 2010). Interestingly, HIF-2α S345 resides within the PAC motif which is needed to maintain correct PAS domain folding (Ponting & Aravind, 1997). Given the transcriptional analysis results (Figures 1-2), the effect of S345 HIF-2α phosphorylation on its interaction with HIF-1β was next investigated using co-immunoprecipitation (co-IP). In HeLa cells, both wild-type and S345A HIF-2α were able to bind HIF-1β, although there was a marked reduction in the binding of S345A compared to wild-type protein. In contrast, interaction of the phosphomimetic S345D HIF-2α variant with HIF-1β was essentially abolished (Figure 3A-B).

**Figure 3.**
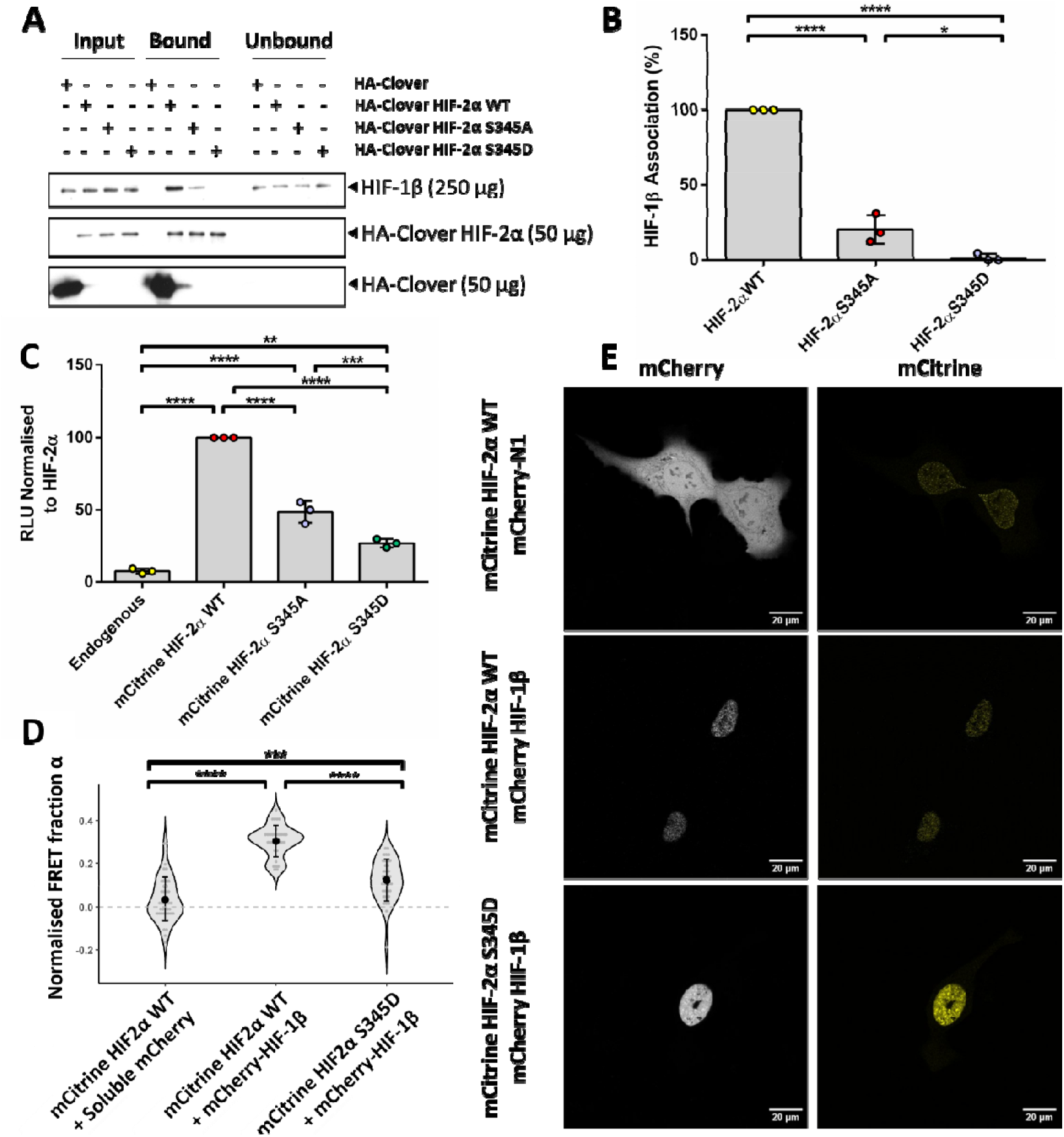
HIF-2α S345D results in reduced HIF-1β binding. (**A** and **B**) Cell lysates from HeLa cells transiently overexpressing HA-Clover or HA-Clover HIF-2α wild-type (WT)/S345A/S345D were processed by GFP Trap co-IP and probed for HIF-1β association. (**A**) Co-IP Western blot probing for Clover and HIF-2α as a loading control. (**B**) HIF-1β band intensity from each condition were quantified and normalised against levels from HA-Clover HIF-2α WT. (**C**) U2OS cells were transiently transfected for low level expression of mCitrine HIF-2α WT/S345A/S345D alongside a HRE luciferase reporter and exposed to 21 % O_2_. HIF activity was normalised to protein concentration, followed by normalisation to the mCitrine-HIF-2α WT condition. (**D**-**E**) FLIM measurements for the HIF-2α WT/S345D interaction with HIF-1β were assessed using U2OS cells transiently transfected with mCitrine HIF-2α WT/S345D alongside either mCherry-N1 or mCherry-HIF-1β and exposed to 21 % O_2_. (**D**) FRET fraction α data for ≥30 cells per condition were obtained. (**E**) Live cell images were taken with a 40x objective on a Leica TCS SP8 confocal microscope. A minimum of 10 cells were imaged per biological replicate (*n* = 3). Statistical differences were assessed using Tukey HSD One-Factor ANOVA with a Bonferroni correction. p > 0.05 is non-significant, p < 0.5 = * and p <0.005 = **.

Previous works have demonstrated how HIF dimer formation can be quantifiably assessed in a live-cell context by FRET-FLIM microscopy (Schutzhold *et al*, 2018). To confirm the observation that HIF-2α S345D impairs HIF-1β association in an *in cellulo* context, a FLIM-FRET approach was employed. U2OS cells were used in this context due to their large, easily identifiable nuclei, which is advantageous when investigating nuclear protein-protein interactions via live-cell microscopy. To this end, mCitrine-tagged HIF-2α wild-type, S345A and S345D constructs were generated. To confirm that the new constructs were not altering the function of HIF-2α, a HRE luciferase assay was performed confirming similar changes in activity of these phosphosite variants compared to wild-type as previously observed (Figure 3C). A striking, and significant, reduction in FRET signal was observed for mCitrine-HIF-2α S345D in combination with mCherry-HIF-1β when compared to HIF-2α wild-type in combination with mCherry-HIF-1β (Figure 3D). Together, these data strongly suggest that HIF-2α S345 phosphorylation impairs HIF transcriptional activity by abrogating HIF-2α interaction with HIF-1β.

The localisation of HIF-2α S345 within the PAC motif (Figure 1A), and its complete conservation across vertebrates suggests that this residue may play a critical role in the correct functionality of the PAC motif, thus regulating HIF dimerisation. Recent advances in protein structure prediction, particularly using tools such as AlphaFold 3 (AF3) and Boltz-1, have revolutionised the study of intermolecular interactions and PTMs, enabling the potential impact of PTM-directed structural changes to be evaluated (Abramson *et al*, 2024; Medvedev *et al*, 2025; Wohlwend *et al*, 2024). Utilising ABCFold, a novel, streamlined workflow integrating AlphaFold 3 and Boltz-1 (Elliott *et al*, 2025), the structural impact of both the S345D, S345A mutations and S345 phosphorylation (pS345) within the HIF-2 heterodimer were investigated.

To date, only a crystal structure of the mouse HIF2 dimer has been solved, but the human proteins are very similar ((Wu *et al*, 2015), PDB: 4ZP4). Protein sequences used for investigating the HIF2 dimer were therefore based upon the mouse sequences of this crystal structure. The AF3 models completed gaps in the 4ZP4 crystal structure from regions that were not fully resolved and, as expected, strongly resembled the crystal structure and had high overall pTM and ipTM scores suggesting confident modelling of both individual proteins and the interface (Sup. Table 1-2). In addition, good ipTM scores were identified for all AF3 models, with the lowest being HIF-2α pS345, suggesting that the interface between proteins in each model have been modelled well when compared to the 4ZP4 structure.

In the wild type AF3 model, the S345 side chain occupies a pocket and points towards the centre of the domain making hydrogen bonds with W318 and E346. When compared to the WT, a consistent local shift in the peptide backbone near residue 345 was observed in all mutant variants (Figure 4A). This can easily be rationalised for the S345D mutant and the phosphorylated S345 since these changes increase the size of the side chain so that it no longer fits tightly in the pocket. The Asp side chain is partially accommodated since the backbone has shifted but makes one rather than two hydrogen bonds (Figure 4B). The even larger phospho group of the pS345 structure is not accommodated at all and makes no hydrogen bonds. Interestingly, although the new side chain of the S345A mutant is smaller, a similar backbone shift is observed, possibly explained by the lack of the tethering hydrogen bonds observed in the wild-type structure. Moreover, the complete loss of hydrogen bonding in the pS345 and S345A structures correlates with additional loss of an intramolecular salt bridge between E346 and K242 of HIF-2α (Figure 4B). It has been argued that AlphaFold models are largely unsuitable for interpretation of single mutations since the coordinates of wild-type and mutant proteins typically differ little (Buel & Walters, 2022). The observation here of a consistent shift of the backbone near S345 upon three diverse alterations is notable. Beyond this, AlphaFold revealed two additional hydrogen bonds between the PAC motif of wild-type HIF-2α and HIF-1β, which are lost in all mutant variants. One of these interactions occurs between E348 of HIF-2α and G294 of HIF-1β, another highly conserved residue downstream of the S345 phosphosite (Sup. Dataset 1). Interestingly, these hydrogen bonds involve residues of HIF-1β that appear to be highly flexible, having not been resolved in the 4ZP4 crystal structure.

**Figure 4.**
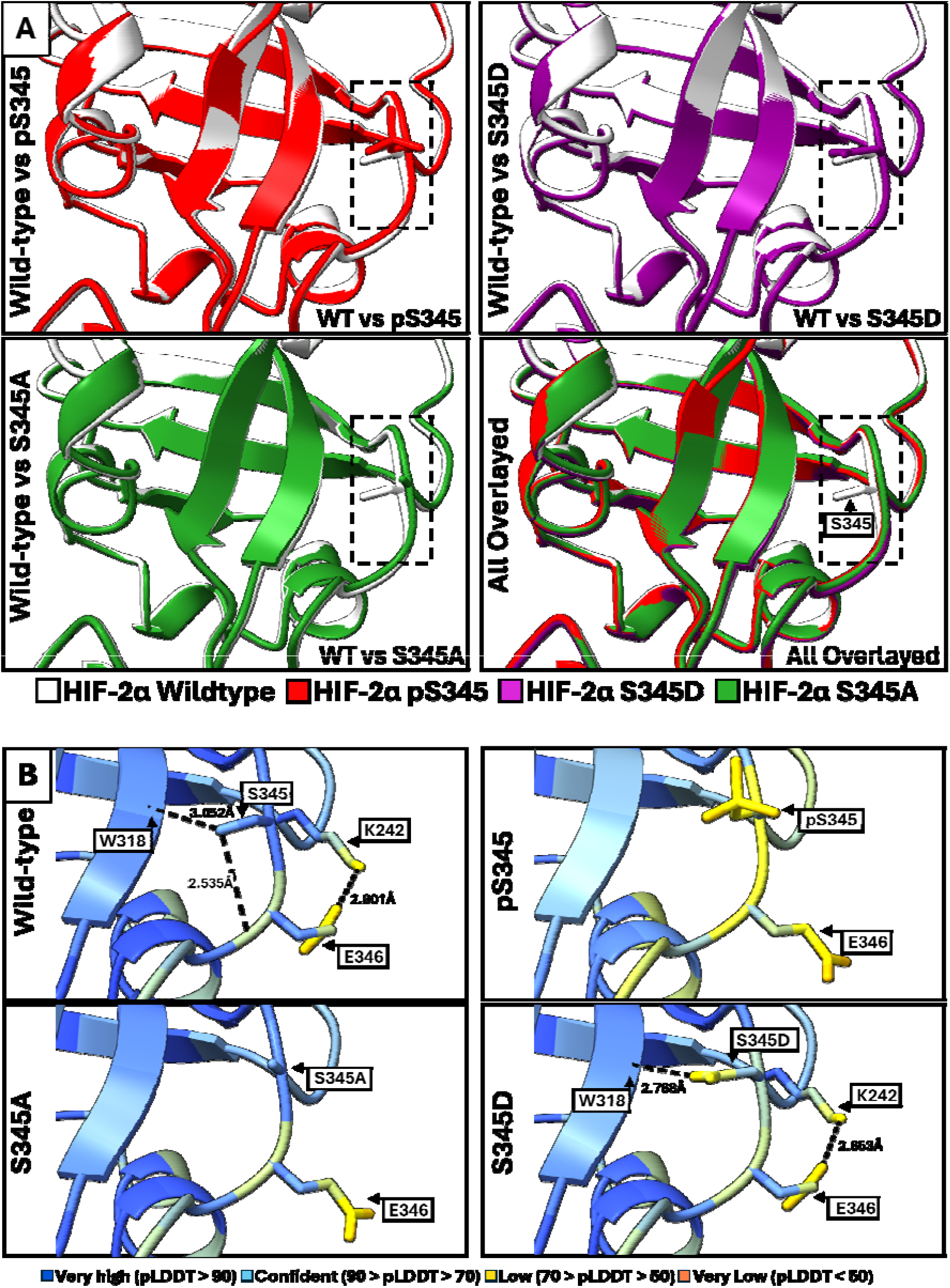
AlphaFold3 modelling of either wild-type, S345A, S345D or pS345 HIF-2^α^ complexed with HIF-1β. Protein sequences utilised for AF3 modelling were based upon the experimentally derived mouse HIF-2α/HIF-1β complex from Wu *et al*., 2015 (PDB: 4ZP4) and visualised using ChimeraX. HIF-1β is hidden from view in this representation. (**A**) The HIF-2α wild-type (WT) model were superimposed with pS345, S345D, S345A separately and all together. A dashed box was used to highlight the location of residue 345, shown in a stick representation, and the local peptide backbone containing it and coloured according to each respective HIF-2α variant. (**B**) HIF-2α variant were also coloured according to pLDDT score, with hydrogen bonding by residues 345 and 346, calculated by ChimeraX, being denoted with bond distances in Angstroms (Å).

Previous work has suggested that the deleterious effect of some mutants can be inferred in AlphaFold models not (principally) from changes in coordinates, but rather from changes in local pLDDT confidence scores. The change in pLDDT upon mutation correlates negatively, albeit weakly, with measured ddG of folding (Zhang *et al*, 2021) and it was observed that the pLDDT of neighbouring residues is affected in the same direction. Relating to the inference that AlphaFold has implicitly learned an energy function (Roney & Ovchinnikov, 2022), the logic is that the placing of a residue in unsuitable, energetically unfavourable environment can be read out from elevated pLDDT scores, both for the residue in question and for others in the vicinity as favourable native packing interactions are disrupted. This feature is strikingly evident in the pS345 and S345D structures (Figure 4B). For example, the mean pLDDT for residue S345 is 74.3 and 83.4 in the pS345 and S345D structures, respectively, compared to 92.3 for wild type HIF-2α.

Taken together, the mutations presented here appear to destabilise the loop containing the S345 residue. In addition to this, the total loss of intra-residue hydrogen bonding proximal to pS345, results in disruption of intermolecular forces of attraction between interfacing residues of HIF-2α and HIF-1β. The increased flexibility of this loop, along with weakened interactions between HIF-2α and HIF-1β, imposes a thermodynamic penalty on interface formation due to reduced entropy, thus possibly explaining the reduced binding between these proteins.

### HIF-2α S345 phosphorylation impairs HIF-1β binding without significantly disrupting other interactions

The results obtained thus far revealed that HIF-2α S345D impairs the fundamental HIF-1β protein interaction required for its transcriptional function. Thus, it was important to determine the extent to which S345D impairs HIF-2α interactions. To this end, a comparative label-free quantitative (LFQ) proteomics analysis was undertaken, analysing the binding partners of wild-type and S345D HIF-2α following immunoprecipitation from HeLa cells using a GFP-Trap approach. Relative quantification revealed similar levels of HIF-2α between wild-type and S345D mutant recovered by this approach (Sup. Dataset 2). Following removal of those proteins observed with the Clover vector control, HIF-2α wild-type was found to interact with 772 proteins, while HIF-2α S345D interacted with 825 (Sup. Dataset 2). Interestingly, over 80% of the interacting proteins were common between these two forms of HIF-2α, with 17% (158) and 11% (105) of the identified protein binding partners for either S345D or wild-type HIF-2α respectively being variant specific (Figure 5A-B). The relative abundance and the proportion of common binding partners indicates that the S345D mutation is not highly disruptive to HIF-2α function, supporting our previous findings presented here.

**Figure 5.**
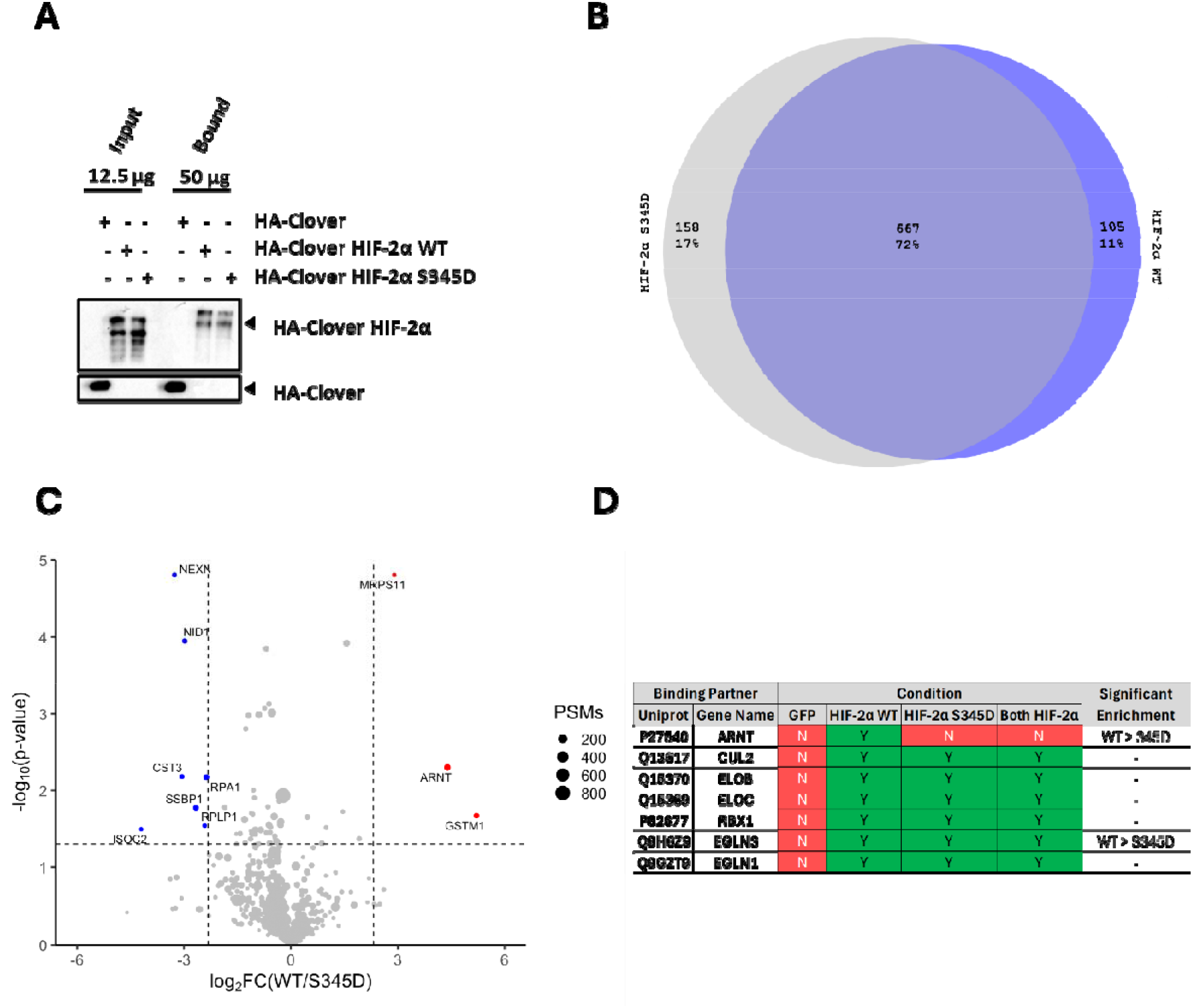
Unbiased binding partner screen of HIF-2α wild-type and S345D protein binding partners using mass spectrometry. (**A**) Western blot validation of GFP-Trap co-IP mass spectrometry binding partners of HIF2α WT/S345D. Cell lysates from HeLa cells transiently overexpressing HA-Clover or HA-Clover HIF-2α WT/S345D were processed for GFP-Trap co-IP Western blot and probed for various binding partners identified by co-IP mass spectrometry. (**B**) Proportion of unique and shared binding partners between HIF-2α wild-type (WT) and S345D when overexpressed in HeLa cells. Venn Diagram of shared and unique protein binding partners of HA-Clover HIF-2α WT/S345D in HeLa cells, displaying the number of interactors and the relative percentage of unique and shared interactors post background subtraction of HA-Clover-only binding partners. Proteins observed in two-of-three biological replicates were maintained in the dataset. A 1 % FDR cut-off was applied after the background subtraction. Circles reflecting each condition are scaled relative to the number of binding partners in each section. (**C**) Volcano plot depicting ≥5-fold significant enriched binding partners of HIF-2α WT and S345D by label-free quantification using complete subtraction of binding partners observed in HA-Clover-Only HeLa cells. Volcano plot displays -log10 (p-value) against log2 abundance fold change to HIF-2α WT/S345D. Each replicate was normalised to HA-Clover HIF-2α abundance. Negative log2 fold change represents enrichment to HIF-2α S345D, and positive values to HIF-2α WT. The total number of MS/MS events is represented by dot size across all replicates. Grey dots represent p-value >0.05 as well as binding partners with p-value < 0.05 but <5-fold significant enrichment. Binding partners with ≥5-fold significant enrichment are labelled with their gene names. Significantly enriched binding partners for HIF-2α WT and S345D are coloured as red and blue, respectively. (**D**) Canonical HIF-2α binding partners observed by GFP-Trap co-IP mass spectrometry. Fundamental binding partners involved in regulating HIF-2α protein stability and activity are organised depending on what experimental condition they were observed in by GFP-Trap co-IP mass spectrometry. Binding partners are grouped depending on their functional roles in HIF regulation. Binding partners that were significantly enriched to either HIF-2α WT or S345D have been annotated. The experiment was carried out *n* = 3.

Importantly, in line with the *in vitro* and *in cellulo* data, HIF-1β was not detected in the HIF-2α S345D interactome but was observed in all replicates of the HIF-2α wild-type interactome (Sup. Dataset 2 and Figure 5C). Indeed, HIF-1β (ARNT) was one of the most strikingly enriched protein binding partners of HIF-2α wild-type compared with S345D (Figure 5C). Interestingly, PHD3, gene symbol *EGLN3*, is likewise enriched as a binding partner of HIF-2α wild-type, although it is unclear as to whether this is an artifact of its increased protein expression levels as evidenced previously by Western blot and RT-qPCR. Beyond PHD3, no other member of the canonical PHD/VHL axis appeared to be differentially altered as a result of the S345 mutation, with both proteins interacting with VHL, Cullin-2, Elongin-B, Elongin-C, RBX-1 and PHD2 (gene name EGLN1) (Figure 5C).

To help interpret the ramifications of S345D on HIF-2α function based upon changes to its interactome, gene ontology (GO) analysis was performed (Sup. Dataset 3). Interestingly, while the unique interactome of HIF-2α wild-type appears to enrich for proteins involved in mitochondrial processes, with the only term with significant FDR (FDR <0.5) being mitochondrial inner membrane (Figure 6A), unique S345D interacting proteins were enriched for multiple terms involved in RNA binding and processing (pre-mRNA splicing), as well as cytoskeletal rearrangement (Figure 6B). While our understanding for the implications of HIF-2 on cytoskeletal regulation is limited, a role for HIF-2 in the formation of a translation initiation complex through direct binding has previously been demonstrated (Uniacke *et al*, 2012). While neither of the two notable proteins that bind HIF-2α, RBM4 and eIF4E2, were identified as interactors of either wild-type or S345D, many other eukaryotic translation initiation factors were identified, with EIF1, EIF2S1, EIF3M and EIF4G2 appearing exclusive to HIF-2α S345D. Beyond this, the enrichment of terms related to pre-mRNA splicing, including multiple components of the spliceosome complex, in the HIF-2α S345D interactome is notable. Together, this data suggests that S345D has broader functional ramifications beyond the immediate notable effect on gene transcription.

**Figure 6.**
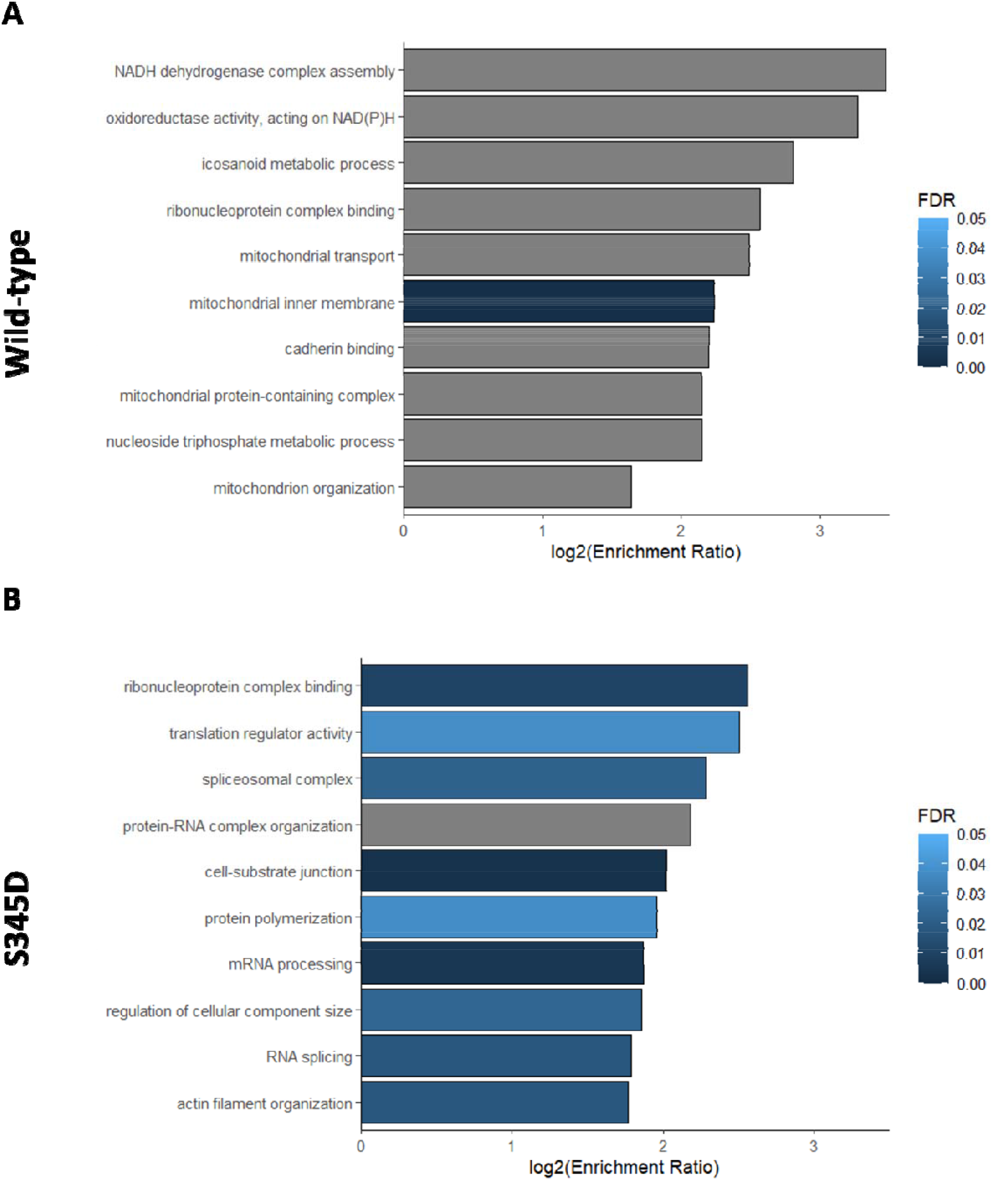
Gene Ontology term enrichment for HIF-2α wild-type and S345D. GO enrichment analysis was performed using WebGestalt against a whole-genome background. The most strongly enriched GO terms of (**A**) HIF-2α wild-type and (**B**) HIF-2α S345D are organised according to their respective log_2_ enrichment ratio. Terms are coloured according to their respective false discovery rate (FDR), with terms denoted in grey not reaching a minimum threshold of FDR ≤ 0.05.

HIF-2α was shown to display a speckle localisation (Taylor *et al*, 2016) and to contain two putative speckle-targeting motifs between residues 450-478 and 771-799 (Alexander *et al*, 2021; Alexander *et al*, 2025). Given that shifts in nuclear speckles appear to correlate with shifts in HIF-2α functional programs, and the substantial enrichment of GO terms relating to RNA binding and processing for S345D, confocal microscopy was employed to evaluated changes in HIF-2α nuclear speckling in the context of S345D (Figure 7). While HIF-2α S345D significantly reduced the number of speckles per cell when compared to wild-type, these speckles displayed significantly enhanced particle size (Figure 7A-C). Together this suggests that HIF-2α S345 phosphorylation may concentrate HIF-2α specific subnuclear foci. Several studies have identified the protein composition of nuclear speckles (Alexander *et al*., 2025; Dopie *et al*, 2020). Comparing the interactome of HIF-2α S345D and HIF-2α WT to that of speckle composition, revealed elevated binding of HIF-2α S345 to over 20 speckle proteins, while HIF-2α WT only exhibited enhanced binding to 4 (other) speckle proteins (Sup. Table 3), suggesting that HIF-2α S345D enhances functions associated with nuclear speckles.

**Figure 7.**
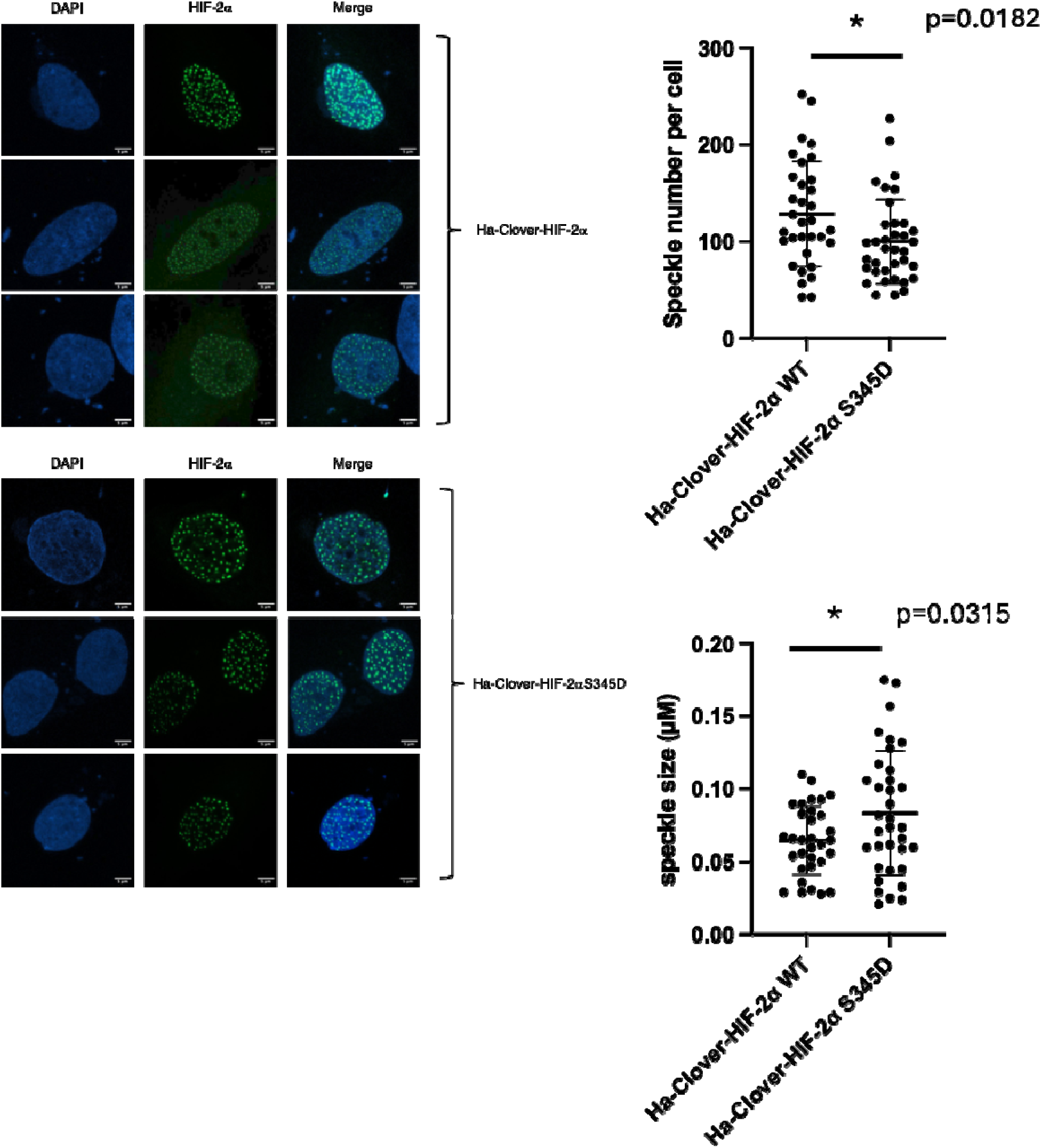
Distinct speckle localisation of HIF-2α S345D. (A) Subnuclear localisation of HA-Clover HIF-2α wild-type (WT) and S345D were assessed by fixed cell 50x confocal microscopy. Representative images are shown with scale bar representing 5µM (B) Quantification of HIF-2α speckle number per cell and (C) speckle size using Image J analysis of confocal images from three independent experiments with each condition having a minimum of 30 cells imaged per biological replicate. Graphs represent all data points with mean and standard deviation also shown. Anova statistical test was calculated to determine statistical significance between conditions (p* <0.05).

Nuclear speckles are hub of RNA processing and metabolism (Abramson *et al*., 2024; Ilik & Aktas, 2022). Given the results obtained by our gene expression analysis, combined with the proteomic and imaging approaches, we next investigated if HIF-2α S345D had enhanced RNA binding properties. As mentioned, HIF-2α has previously been shown to associate with proteins related to mRNA binding and, for example, interact with EGFR mRNA via the 3’-UTR, in a HIF-1β independent manner (Uniacke *et al*., 2012).

RNA-IPs were carried out for HIF-2α wild-type and S345D and probed for their respective interaction with 3’-UTR of EGFR. To this end, two distinct primer sets were generated against the 3’-UTR of EGFR (Figure 8) based upon the work carried out by Uniacke *et al*. (Uniacke *et al*., 2012). Expression of either Clover-only, HIF-2α wild-type or S345D did not increase EGFR transcript levels, indicating that any specific mRNA binding is not artificially enriched by elevated transcript levels in the cell (Figure 8A-B). Analysis of the results obtained using the primer targeting EGFR 3’ UTR nucleotides 4295-4657 highlighted a significant increase in specific mRNA binding by HIF-2α wild-type and S345D alike, but with a further significant increase from S345D when compared to wild-type (Figure 8D). No significant change in mRNA binding was observed with the second primer set targeting nucleotides 4657-4861 when comparing GFP to HIF-2α wild-type, despite a similar trend being observed (Figure 8E). However, a significant increase in specific mRNA binding was observed when comparing GFP to S345D. Binding to RPL13A was used as a negative control and revealed no differences between Clover-only and any of the HIF-2α constructs used (Figure 8C). This observation of variable UTR binding dependent on primer set was likewise evident during the Uniacke *et al*. (Uniacke *et al*., 2012) investigation.

**Figure 8.**
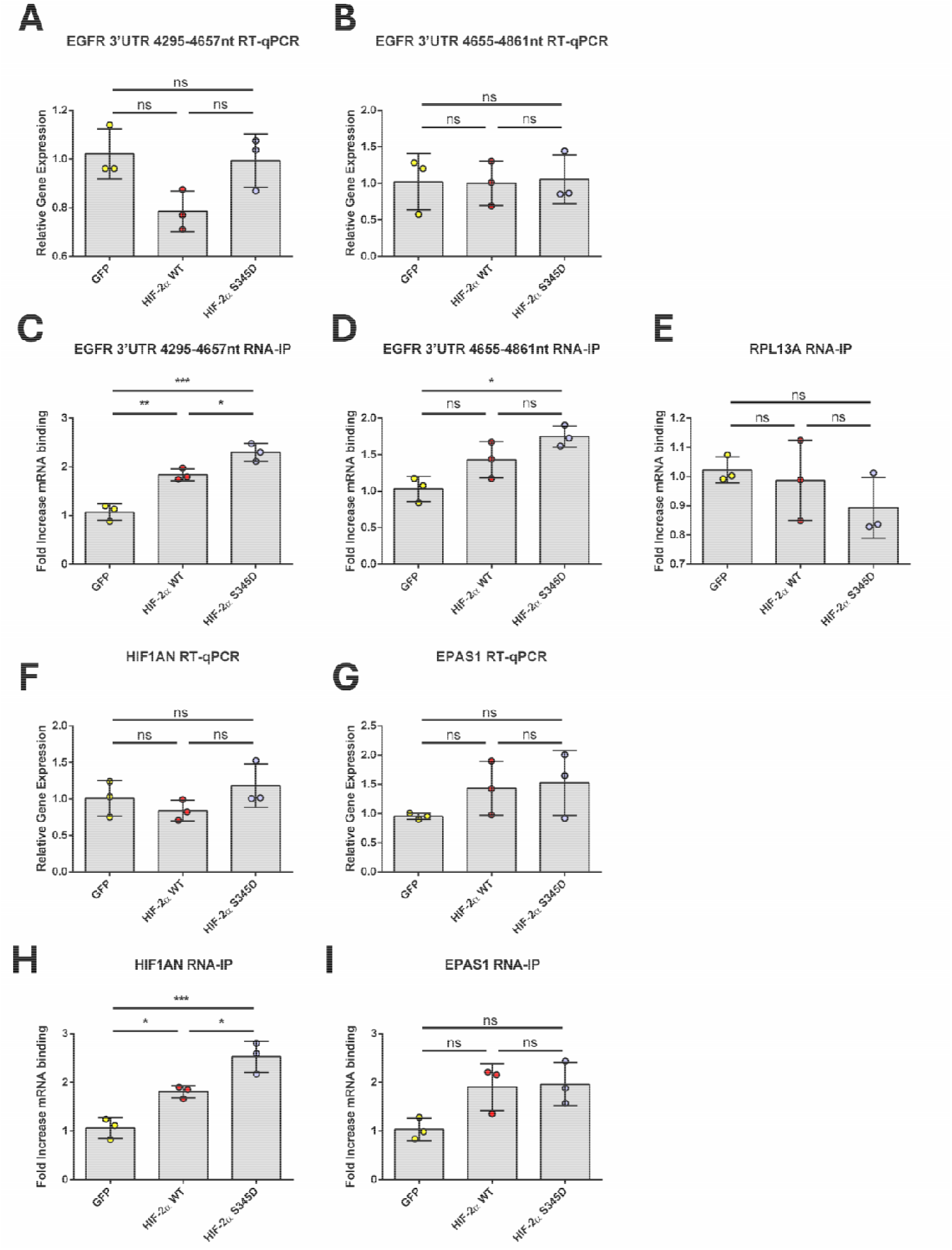
GFP-Trap RNA-IP for HA-Clover HIF-2α wild-type and S345D. Gene expression levels of EGFR (**A** and **B**), HIF1AN (F) and EPAS1 (G) mRNA was quantified by RT-qPCR following overexpression of either HA-Clover-only or HA-Clover HIF-2α wild-type (WT)/S345D, normalised against RP13A mRNA levels. Two primer sets against the 3’ UTR of EGFR mRNA were used (targeting nucleotides 4657-4861 and 4295-4657). (**C, D. E, H and I**) HeLa transiently overexpressing HA-Clover-only or HA-Clover HIF-2α WT/S345D were crosslinked to preserve interactions with protein/DNA/RNA and processed by GFP-Trap. Crosslinking was then reversed. RT-qPCR was carried out using the two primer sets against the EGFR 3’ UTR, HIF1AN 3’ UTR, EPAS1 3’ UTR and RPL13A as a loading control. One-way Anova Tukey post-hoc test was performed to determine significance (*P < 0.05; ns, not significant). *n* = 3.

Beyond EGFR, Uniacke *et al*. (Uniacke *et al*., 2012) identified a wide range of HIF-2α mRNA interactors in their RNA-IP sequencing approach, including HIF1AN and EPAS1, translated to FIH and HIF-2α, respectively. Given that HIF activity is very tightly controlled by multiple intricate feedback loops, the implications of HIF-2α interacting with its own mRNA as well as FIH, a well-established negative regulator of HIF activity, is interesting (Lando *et al*., 2002; Mahon *et al*., 2001). To further explore the role of S345D in HIF-2α specific mRNA binding, both HIF1AN and EPAS1 mRNAs were investigated by RNA-IP. Once again, no increase in HIF1AN or EPAS1 endogenous transcript levels were detected when HeLa cells were exogenously expressing either Clover-only, HIF-2α wild-type or S345D (Figure 8F-G). In line with HIF-2α specific binding of EGFR transcripts, compared to the Clover control a significant increase in HIF1AN specific mRNA binding was observed for HIF-2α wild-type and S345D, but with a further significant increase from S345D when compared to wild-type (Figure 8H). Contrasting this, there was no significant change in the binding of either HIF-2α wild-type or S345D to EPAS1 transcripts via the 3’ UTR (Figure 8I). Therefore, this suggests a role for HIF-2α S345 phosphorylation in controlling specific mRNA binding.

## Discussion

The data presented here suggests that phosphorylation of S345 in HIF-2α serves as a switch to enhance RNA binding and reduce its activity as a transcription factor which requires binding to HIF-1β. Although, phosphorylation has been shown to change HIF-1α localisation (Gkotinakou *et al*., 2019; Mylonis *et al*., 2008; Mylonis *et al*., 2006; Pangou *et al*, 2016), in the case of S345 of HIF-2α no changes in expression levels or overall nuclear localisation were detected. Furthermore, most of HIF-2α binding partners were preserved, suggesting that S345D mutation is not causing drastic structural or localisation changes to the HIF-2α protein. This suggests a finer tuning of the function following this phosphomimetic modification. Phosphorylation of S345 of HIF-2α was originally identified following 4h hypoxia exposure (Daly *et al*., 2021). At present we have no information on the dynamics of stoichiometry of this modification. It is possible that this phosphorylation represents a temporal component of regulation, shifting the pool of HIF-1β to HIF-1α during acute hypoxia. Interestingly, although the consensus motif around this HIF-2α site is conserved in HIF-1α, modification of HIF-1α was not detected in the original study, even though this region of the protein was observed (Daly *et al*., 2021). It remains a possibility though that equivalent HIF-1α phosphorylation may occur with different dynamics or during more chronic hypoxic events. Additional research is therefore needed to address these possibilities.

### Role of HIF-2α in mRNA processing and translation

Protein synthesis, an inherently energy-intensive process, undergoes extensive regulation during hypoxia. Under severe conditions, ATP demand for protein synthesis is decreased to 7 % of normoxic levels, correlating with a decline in protein translation (Hochachka *et al*, 1996). Several signalling cascades, including mTORC1 and PKR-like ER kinase (PERK), are implicated in this downregulation (Reviewed by (Ivanova *et al*, 2019)). mTORC1 impedes translation initiation by suppressing eukaryotic translation initiation factor 4F (eIF4F) complex recruitment to 5’ cap of mRNA transcripts. PERK is central to the unfolded protein response (UPR), a molecular response to severe hypoxia/anoxia (<0.2 % O_2_) that limits energy consumption as well as global protein synthesis (Koritzinsky *et al*, 2006). Activated PERK phosphorylates eIF2α, which in turn becomes a dominant negative inhibitor of eIF2B, thereby suppressing both translational initiation and global protein synthesis (Sonenberg & Hinnebusch, 2009; Wek & Cavener, 2007). Interestingly, mTOR and eIF2α were identified as novel HIF-2α binding partner by MS during this investigation. While mTOR was not identified as a significantly enriched binding partner by LFQ to either HIF-2α wild-type or S345D, RICTOR was. mTOR and RICTOR are two fundamental components of mTORC2, which is known to regulate HIF-2α on transcriptional and translational levels (Mohlin *et al*, 2015; Nayak *et al*, 2013). While mTORC1 is known to regulate HIF-1α by binding to a mTOR signalling motif immediately downstream of the PAS-A domain to regulate CBP/p300 binding, the consequences of an interaction between mTORC2 and HIF-2α remains unknown.

Our interactome proteomic data suggests that S345 of HIF-2α is involved in RNA binding. We were able to detect increased binding to two HIF-2α RNA targets, EGFR and HIF1AN, described in Uniacke *et al*. (Uniacke *et al*., 2012). In addition, the implications of HIF-2α binding to HIF1AN mRNA, a canonical regulator of HIF-α activity are not yet understood. This suggests that perhaps phosphorylation of S345 of HIF-2α is shifting HIF-2α from binding DNA with its binding partner HIF-1β to binding mRNA with RNA binding proteins, as well as proteins involved in translation and RNA processing (Figure 9). Ideally, an unbiased analysis of HIF-2α binding to RNAs would be necessary to fully determine the extent of RNA control by this HIF protein. This other aspect of HIF-2α biology is also an important one: as more patients will be treated with Belzutifan (Deeks, 2021), separating HIF-2α from HIF-1β will not only lead to reduced transcriptional activity, but could result in increased RNA binding, for the fraction of HIF-2α that escapes degradation and/or is phosphorylated at S345.

**Figure 9.**
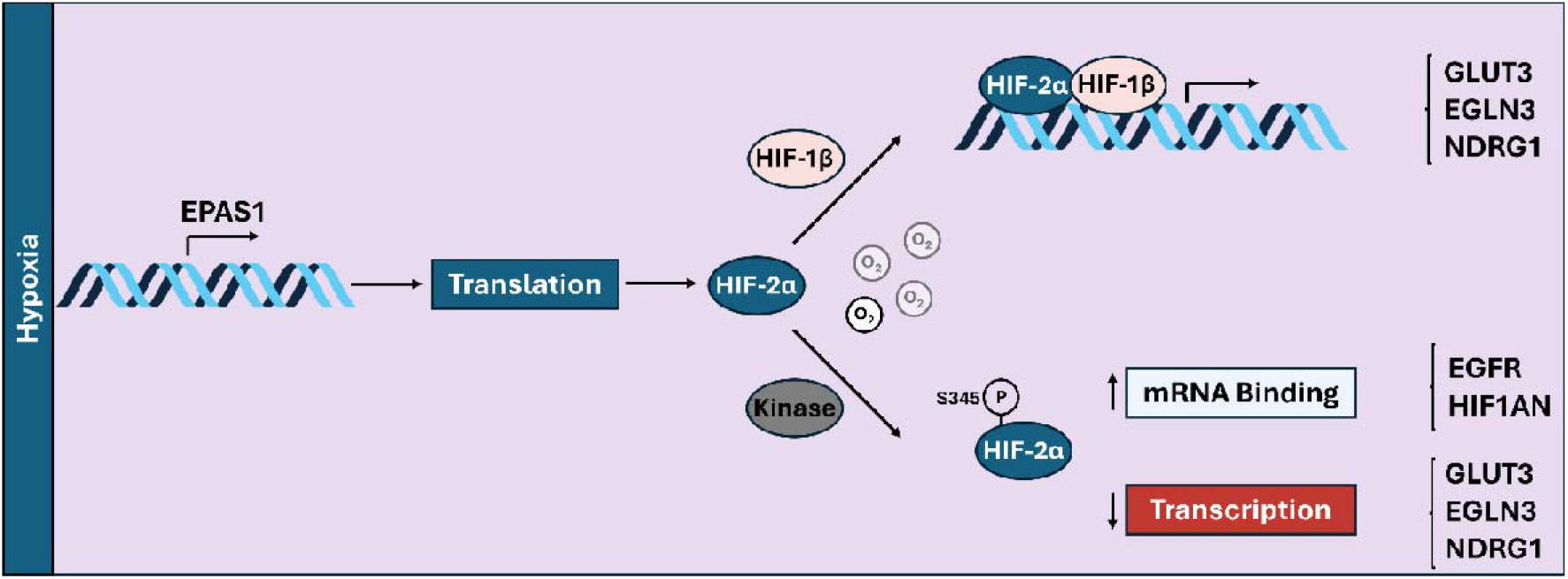
Proposed functional consequences of HIF-2α S345 phosphorylation. Following hypoxic stabilisation of HIF-2α, HIF-2α can be phosphorylated as S345 to abolished HIF-1β heterodimerisation, impairing HIF-2 dependent gene expression. Despite the impairing transcriptional activity, HIF-2α S345 phosphorylation increases binding of specific mRNA.

Beyond translational regulation, GO analysis of the unique interactomes of HIF-2α wild-type and the S345D mutant revealed an enrichment for terms related to mRNA processing, including ribonucleoprotein complexes and spliceosome components, in the S345D dataset. The spliceosome, composed of ribonucleoprotein complexes, coordinates pre-mRNA splicing (Sharp, 1994; Zhou *et al*, 2002). Interestingly, ribonucleoproteins, much like HIF-2α itself, are known to assemble into nuclear speckles (Hall *et al*, 2006; Spector & Lamond, 2011; Taylor *et al*., 2016). HIF-2α was recently suggested to contain two internal speckle-targeting motifs (STMs), and that alterations in its speckling signature were linked to changes in functional programmes and the subset of target genes preferentially transcribed by HIF-2α (Alexander *et al*., 2025). Nuclear speckling has also been implicated in cancers such as clear cell renal cell carcinoma (ccRCC), where functional changes in HIF-2α mediated by speckle dynamics have been proposed to underlie differences in patient survival and may influence tumour responsiveness to specific therapies (Regan-Fendt & Izumi, 2024).

Despite these advances, the mechanisms driving transitions between speckle states, and the biological significance of differential speckling in normal physiology, remain unclear. In this context, the observation that the HIF-2α S345D phosphomimetic markedly alters its nuclear speckling is particularly intriguing. The reduction in foci number, coupled with an increase in speckle size, suggests that phosphorylation at S345 promotes the concentration of HIF-2α into a more restricted number of enlarged nuclear puncta.

Whereas Alexander et al. (Alexander *et al*., 2025) associated changes in HIF-2α speckling with selective induction of HIF target genes, the S345D mutant is transcriptionally inactive and instead shows enrichment of GO terms related to RNA binding and processing. The enrichment of mRNA-processing proteins among HIF-2α S345D interactors raises the possibility that non-dimerised HIF-2α may play a previously unrecognized role in regulating RNA, extending beyond translational control (Uniacke *et al*., 2012) to include other aspects RNA such as splicing. If so, this would suggest a novel mechanism of hypoxia-mediated cellular regulation—one that not only controls which genes are expressed but also influences how those genes are spliced. Such regulation could potentially affect mRNAs outside the canonical HIF transcriptional program, thereby shaping the proteome by directing the production of specific splice variants. Further work is required to establish whether this novel mode of regulation is coupled to the HIF-2α S345D-mediated increase in mRNA binding shown by this investigation.

### Limitation of the study

Our study mostly used phospho-mimetic/null mutation of HIF-2α at S345. We were unable to identify the upstream kinase responsible for deposition of site-specific phosphorylation. Knowing the identity of the kinase would allow more physiological regulatory mechanisms to be investigated. Despite several kinases fitting the consensus for HIF-2α S345, many of the prediction tools, including GPS 6.0, GPS 5.0 and PhosphoSitePlus, suggest kinases with very generic consensus motifs which reduce the likelihood of them being true regulators, including casein kinase 1 (CK1) (Chen *et al*, 2023; Johnson *et al*, 2023; Wang *et al*, 2020). To date, CK1 is recognised to phosphorylate HIF-1α at S247 to impair HIF-1β binding (Kalousi *et al*., 2010). However, with respect to HIF-2α, CK1δ is suggested to phosphorylate S383 and T528 to promote HIF-2α nuclear retention, with no apparent effect on HIF-1β binding in HeLa cells (Pangou *et al*., 2016). An alternative predicted kinase includes CK2α which, much like CK1, has a very generic consensus motif. Gradin *et al*. (Gradin *et al*, 2002) have previously suggested CK2α to be a positive regulator of HIF-2α transcriptional activity. It is likely that an unbiased approach for upstream regulator identification is necessary, such as a kinase screen against a purified peptide consisting of the S345 consensus.

Although, using candidate approaches and generic luciferase reporter essays, HIF-2α S345D was inactive, it is still possible that it can bind another partner to control gene transcription away from HIF-1β binding. From the proteomic analysis we could only detect STAT1 as another transcription factor with enhanced binding to HIF-2α S345D. However, it is unknown if these two proteins are able to productively control transcription and it would be interesting to investigate this further. Reconstitution of HIF-2α null cells with HIF-2α wild type and S345D combined with RNA-sequencing would answer this question.

## Experimental Methods

### Cell culture, hypoxia treatment and lysis

HeLa cervical cancer cells, human osteosarcoma U2OS and HEK-293T human embryonic kidney cells were purchased from the A.T.C.C. All cells were grown in Dulbecco’s modified Eagle’s medium with 10% (v/v) foetal bovine serum and 1% MEM Non-Essential Amino Acids Solution at 37 °C, 5 % CO_2_. Hypoxic incubations were carried by exposing cells to 1% oxygen, 37 °C, 5 % CO_2_ in an InVivo hypoxia workstation (Ruskin).

Cells were lysed two days post-transfection. After removal of the media, cells were washed two times with PBS. For immunoblotting and proteomic analysis, cells were harvested in lysis buffer (50 mM Tris (pH 8.0), 120 mM NaCl, 5 mM EDTA, 0.5% (v/v) NP-40, 1 Pierce Protease Inhibitor Mini Tablet (ThermoFisher Scientific). Lysates were rotated end-over-end for 10 minutes at 4 °C followed by centrifugation at ≥15,000g for 10 minutes at 4 °C. Supernatants were then collected and quantified by Bradford Assay (BioRad).

### Site-directed Mutagenesis

Plasmid manipulation, including site-directed mutagenesis (SDM), was carried out using the Takara Bio In-Fusion Cloning kit according to the manufacturer’s protocol. All primers were designed using the Takara Bio primer design tool. Custom desalted oligonucleotide primers for cloning were ordered from Sigma Aldrich, and were resuspended to 100 μM in MilliQ water, vortexed and frozen at-20 °C. Primers were diluted to 5 μM stocks for experimental use. The following primers were designed and used for SDM: HIF-2α S345A, 5’-CGTCCTGGCTGAGATTGAGAAGAATGACGTGGTGT-3’ (forward) and 5’-ATCTCAGCCAGGACGTAGTTGACACACATGATG-3’ (reverse) and HIF-2α S345D, 5’-CGTCCTGGATGAGATTGAGAAGAATGACGTGGTGT-3’ (forward) and 5’-ATCTCATCCAGGACGTAGTTGACACACATGATG-3’ (reverse).

### Transient Transfection of plasmid DNA

For immunoblotting and proteomics experiments, mammalian cells were transiently transfected with plasmids using a 1% stock of PEI 40K MAX (Polysciences) (w/v) in phosphate-buffered saline (PBS) (pH 7.5, adjusted with NaOH). Non-supplemented DMEM media was used to dilute DNA to a final concentration of 10 ng μL^-1^. Low and high expression transfections were carried using DNA in a ratio of 1:19 and 1:1 HA-Clover-HIF-α:pcDNA3(-) (Invitrogen #V79520), respectively. Media was changed 16 hours post-transfection. HA-Clover, HA-Clover HIF-2α, HA-Clover HIF-2α S345A and HA-Clover HIF-2α S345D are available via Addgene (163366, and 163371, 163372 and 163373, respectively). mCitrine-C1, mCitrine-HIF-2α, mCherry-C1 and mCherry-HIF-1β and mCitrine-mCherry fusion were gifted from Prof. Joachim Fandrey. mCitrine-HIF-2α S345A and mCitrine-HIF-2α S345D were generated by site-directed mutagenesis using the In-Fusion cloning kit (Takara Bio).

For U2OS cells cultured for FLIM-FRET microscopy, U2OS cells were plated to at least 80 % cell density in 15 μL 8-well glass bottom slides (Ibidi). In a 1.5 mL Eppendorf, 12.5 μL serum-free media was mixed with 300 ng plasmid. The P3000 reagent from the Lipofectamine™ 3000 Transfection Reagent (ThermoFisher Scientific) was added to the serum-free/DNA mix in a ratio of 1Ul P3000:500 ng plasmid DNA. A second Eppendorf was prepared containing 12.5 μL serum-free and 0.75 μL Lipofectamine 3000 reagent. The two tubes were combined, pipetted up/down to mix, and left to incubate for 15 minutes (room temperature) prior to being added dropwise to cells. Media was changed after 6 hours post-transfection.

### Immunoblotting

Primary antibody incubations were carried out overnight (18 hours) at 4°C with HIF-1α (Proteintech #20960-1-AP; 1/1000) and HIF-2α (Bethyl Laboratories #A700-003; 1/1000), β-actin (Abcam #ab8226; 1/10000) and GFP (Roche #11814460001). Secondary antibody incubation was performed for 2 hours at room temperature with anti-rabbit horseradish peroxidase (HRP) (Cell Signaling #7074s; 1/2000) or anti-mouse HRP (Cell Signaling #7076; 1/2000). Blots were developed on film using a Protec ECOMAX X Ray Film Processor.

### Co-immunoprecipitation of HA-Clover HIF-2α for immunoblotting and proteomics

Cells were washed with PBS and lysed in lysis buffer (50 mM tris-HCl (pH 8.0), 120 mM NaCl, 5 mM EDTA, 0.5% (v/v) Nonidet P-40, 1 Pierce Protease Inhibitor Mini Tablet (ThermoFisher Scientific) per 10 mL of buffer). Lysates were rotated for 10 min at 4°C before centrifugation at >15,000g for 10 min at 4°C, following which the clarified supernatant were collected. Supernatants were diluted 2.5-fold in dilution buffer (lysis buffer excluding detergent) to dilute Nonidet P-40 to 0.2% (v/v). For preclearing, 10 μL Bab-20 beads (CHROMOTEK) per sample were washed in 90 μL dilution buffer, centrifuged at 1000 g, and then supernatants discarded. This step was carried out three times. Washed Bab-20 beads were then combined with diluted lysates. Bead/protein suspensions were rotated end-over-end for 1.5 hours (RT). Bead/protein suspensions were then centrifuged at 5000 g for 5 minutes to pellet the Bab-20 beads. 10 μL green fluorescent protein (GFP) -Trap agarose beads (CHROMOTEK) per sample, were equilibrated in an identical manner to the Bab-20 beads. The supernatant was then transferred to fresh tubes and incubated overnight at 4 °C with pre-equilibrated GFP-Trap beads. Following centrifugation at 3000 g for 3 minutes, the unbound supernatant was discarded. Bead/protein suspensions were washed in 200 μL dilution buffer and centrifuged at 3000 g for three minutes. This step was carried out three times. For Western blots, beads were resuspended in 50 μL 5x SLB, heated for 10 minutes at 90 °C, and centrifuged. For mass spectrometry, proteins were eluted with a dedicated buffer (50 mM Tris (pH 8.0), 2 % SDS) and heated at 90 °C with intermittent mixing. If samples were not processed immediately, samples were flash frozen in liquid nitrogen and stored at -80 °C.

### LC-MS/MS

IP preps were diluted to 180 μl in 100 mM ammonium bicarbonate pH 8.0 and reduced and alkylated with dithiothreitol and iodoacetamide, as previously described by (Ferries *et al*, 2017). A 1:1 mixture of magnetic hydrophobic and hydrophilic Sera-mag SpeedBeads (Cytivia) was added at a 2:1 (w/w) ratio of beads/protein and HPLC-grade Acetonitrile (ACN) was added to a final concentration of 80% (v/v) and incubated at 25 °C with shaking (1500 rpm) for 30 min. The supernatant was discarded, and beads were washed (3×) in 200 μL of 100% ACN. Beads were dried by vacuum centrifugation for 10 min and resuspended by water bath sonication for 2 min in 200 μL of 100 mM AmBic and 0.2 μg of Trypsin Gold (Promega) added prior to incubation overnight at 37 °C with 1500 rpm shaking. Beads were removed and supernatant moved into a new tube using a MagRack (Cytivia), followed by Trifluoroacetic acid (TFA) acidification to a final concentration of 0.5%. Samples were incubated at 25 °C with shaking (1500 rpm) for 30 min, followed by 30 min on ice. Samples were centrifuged at 15,000 g for 10 mins and 195 μL moved to a new tube and dried by vacuum centrifugation. All dried peptides were solubilized in 20 µl of 3% (v/v) ACN and 0.1% (v/v) TFA in water, sonicated for 10 min, and centrifuged at 13,000g for 15 min at 4°C before liquid chromatography–mass spectrometry analysis. Samples were separated by reversed-phase HPLC separation using an Ultimate3000 nano system (Dionex) over a 30-minute gradient, as described by (Ferries *et al*., 2017). Data acquisition was performed using a Thermo Orbitrap Fusion Lumos Tribrid mass spectrometer (Thermo Scientific), with higher-energy C-trap dissociation (HCD) fragmentation set at 32% normalized collision energy for 2+ to 5+ charge states using a 3 s cycle time. MS1 spectra were acquired in the Orbitrap (120K resolution at 200 *m/z*) over a range of 350 to 2000 *m/z*, AGC target = 50%, maximum injection time = 50 ms, with an intensity threshold for fragmentation of 1e^4^. MS2 spectra were acquired in the ion trap (rapid mode), AGC target = standard, maximum injection time = dynamic. A dynamic exclusion window of 60 s was applied at a 10 ppm mass tolerance. Data was analysed using Proteome Discoverer 2.4 (Thermo Scientific) in conjunction with MASCOT (Perkins *et al*, 1999); searching the UniProt Human Reviewed database (updated weekly, accessed May 2023) with constant modifications = carbamidomethyl (C), variable modifications = oxidation (M), instrument type = electrospray ionization–Fourier-transform ion cyclotron resonance (ESI-FTICR), MS1 mass tolerance = 10 ppm, MS2 mass tolerance = 0.5 Da. Label free quantification was performed using the Minora feature detector node, calculating the area under the curve for *m/z* values, total protein abundance was determined using the HI3 method (Silva *et al*, 2006). Identifications observed in at least 2 replicates of any conditions were maintained in the dataset and background subtracted. Missing values were imputed using low abundance resampling, T-tests performed and fold changes determined. Data was plotted using a custom R script. Overrepresentation analyses were performed with WebGestalt (Elizarraras *et al*, 2024) against a whole-genome background, using the three Gene Ontology noRedundant databases.

Proteomic data will be available via submission to PRIDE upon publication.

### RNA-Immunoprecipitation

HeLa cells transfected with HA-Clover-Only/HIF2α constructs were resuspended in hypotonic polysome extraction buffer (5 mM Tris pH 7.5, 2.5 mM MgCl2, 1.5 mM KCl, 1 % Triton X-100, 100 mg/mL cycloheximide (CHX), 100 U/mL RNasin) and lysed through the addition of Triton X-100 (0.5 %) and sodium deoxycholate (0.5 %) as described in (Ivanova et al., 2018). Clarified lysates were incubated with GFP-trap beads according to protocols found in 2.7. Bead/protein suspensions were washed with polysome extraction buffer and RNA isolated using the PeqGold RNA isolation kit, as per 68 the manufactures protocol. Copy DNA (cDNA) was prepared using Qiagen Quantanova cDNA synthesis kit according to the manufacturers protocol. Analysis of cDNA samples were carried out as outlined in section 2.9. RPLA13 was used a normalising gene. The following primers were used for RT-qPCR following RNA IP: RPLA13, 5’-CCTGGAGGAGAAGAGGAAAGAGA-3’ (forward), 5’-TTGAGGACCTCTGTGTATTTGTCAA-3’ (reverse), EGFR 3’ UTR 4295-4657nt, 5’-TTCATCCAGGCCCAACTGTG-3’ (forward) 5’-GGGCAGCATACTGAGTTTCA-3’ (reverse), EGFR 3’ UTR 4657-4861nt, 5’-CCCCTGTCTTGCTGTCATGAA-3’ (forward) and 5’-GCCCCAAAGGACCTGATGCAT-3’ (reverse); EPAS1, 5′-TTTGATGTGGAAACGGATGA-3′ (forward) and 5′-GGAACCTGCTCTTGCTGTTC-3′ (reverse). HIF1AN 5′-CATAAAGTCTGCAACATGGAAGGT-3′ (forward) and 5′-ATTTGATGGGTGAGGAATGGGTT-3′ (reverse).

### RT–qPCR

Total RNA was purified from the cells using the peqGOLD Total RNA Kit (VWR Life Sciences) according o the manufacturer’s protocol. The concentration and purity of the extracted RNA were assessed using a Nanodrop 2000c spectrophotometer. RNA was then converted into cDNA using the iScript Reverse Transcription Kit (Bio-Rad) following the recommended protocol. Power-track sybr green master mix (Applied Biosystems) was used to analyse cDNA samples on the Mx3005P RT-qPCR platform with MX3005P 96-well plates (Stratagene/Agilent). Actin was used a normalising gene, and results were analysed using the ΔΔCt method. Primers used for RT-qPCR are listed as follows: Actin 5’-CTGGGAGTGGGTGGAGGC-3’ (forward) and 5’-TCAACTGGTCTCAAGTCAGTG-3’ (reverse), GLUT1 5’-CAAATGACATCCCAGCTTGA-3’ (forward) and 5’-TTGTGTGGTTATCGCTCTCG-3’ (reverse), PHD3 5’-CTTGGCATCCCAATTCTTGT-3’ (forward) and 5’-ATCGACAGGCTGGTCCTCTA-3’ (reverse) and NDRG1 5’-GGAGTCCTTCAACAGTTTGG-3’ (forward) and 5’-CACCATCTCAGGGTTGTTTAG-3’ (reverse).

### Luciferase reporter assays

24 hours post-plating, HeLa/U2OS cells transiently transfected with HRE Luciferase plasmid (and relevant additional plasmids). 16 hours post-transfection, media was replaced, and cells exposed to normoxia or hypoxia (21 % or 1 % O2, respectively) for 24 hours. Cells were then washed with PBS and lysed with 200 μL luciferase lysis buffer (25 mM Tris-phosphate pH 7.5, 1 % (w/v) bovine serum albumin (BSA), 15 % (v/v) glycerol, 8 mM MgCl2, 0.1 mM EDTA, 2 mM DTT, 1 % (v/v) Triton x-100) per well (RT). For each condition, 5 μL lysate was added to a tube, and 50 μL luciferase substrate (Promega) was automatically injected by a Lumat LB 5907 luminometer (EG&G Berthold). Luminescence was 64 measured for 10 seconds without background subtraction. HEK293 cells were prepared similarly, using 400 μL lysis buffer per well.

### Fixed Cell Immunofluorescence Staining and Fixed Cell Microscopy

Cells were seeded onto sterile coverslips in 6-well plates. 24 hours post-seeding, transfections were performed, followed by a media change 16 hours later. After 24 hours of recovery, cells were washed 72 gently with PBS and fixed in ice-cold 100 % methanol for 7 minutes at -20 °C. Methanol was carefully aspirated, and cells were washed three times for 5 minutes each with PBS on a shaker, in the dark using a humidified chamber. Hoechst stain (diluted 1:15000) was pipetted onto each parafilm square (120 μL per square). Coverslips with cells were carefully inverted onto the Hoechst spots for 2 minutes with the lid of the chamber closed. Following incubation, coverslips were briefly dipped in MilliQ water to remove excess stain and gently dried with paper towels. A drop of mounting medium was placed on microscope slides, and coverslips were mounted cell-side down. Excess mounting medium was removed with a tissue, and the edges of the coverslips were sealed with nail varnish. Slides were dried in the dark and stored at -20 °C.

Fixed cells were imaged at 20x magnification (numerical aperture 0.75) on an LSM710 Zeiss Microscope equipped with GFP (488 nm) filters. DAPI fluorescence detection was facilitated by a Picoquant laser controlled through the SymPhoTime64 software package. Representative images and quantification of HA-Clover HIF-2α nuclear accumulation in fixed HeLa were carried out using the Omero.Insight 5.7.1. software package (Allan *et al*, 2012). ROIs were selected for background, nuclear and non-nuclear GFP signal. Nuclear ROIs were generated based upon DAPI signal, and non-nuclear GFP was identified as any GFP signature not co-localising with the DAPI signature. Using ImageJ, ROIs were selected for the Hoechst-stained nucleus to determine the proportion of GFP-tagged HIF-2α in the nucleus and cytoplasm for the HIF-2α wild-type and S345D. In addition, the gross fluorescence levels from cells expressing either the HIF-2α wild-type or S345D were measured.

### Confocal analysis

HeLa Cells were fixed onto coverslips with 3.7% paraformaldehyde in phosphate buffered saline (PBS) for 15 minutes, then rinsed in PBS and permeabilised with 0.5% triton X-100 in PBS for 10 minutes. Prior to staining cells were blocked with 1% donkey serum in PBST (0.05% tween) for 30 minutes). The coverslips were then stained with Hoechst 33342 (1:150000, Invitrogen, H3570) for 2 minutes and mounted onto slides (Vectashield antifade mounting medium, H1000). Images were acquired on the Cell discovery 7 microscope, using the confocal lens at 50 X magnification. Gain and intensity settings of cell discovery 7 microscope: Hoechst 33342 channel, laser intensity 1.5%, gain of 725V; GFP channel, laser intensity 0.8%, gain of 550V. Image quantification was performed in Image J-Fijii particle analysis pipeline.

### FLIM-FRET

U2OS cells were transfected with mCitrine-C1, mCitrine-HIF-2α, mCherry-C1 and mCherry-HIF-1β, mCitrine-mCherry fusion, mCitrine-HIF-2α S345A and mCitrine-HIF-2α S345D as described in Schützhold et al. (2018). Live cell images were taken with a 40x objective on a Leica TCS SP8 confocal microscope with SP FLIM (TCSPC module PicoHarp 300, Picoquant) and analysed as described before (Schutzhold *et al*., 2018).

### Evolutionary Analysis

Evolutionary analysis of HIF-1α and HIF-2α protein vertebrate and invertebrate homologues were performed using HIF-α homologue lists as described in Daly *et al*., 2021. The MUltiple Sequence Comparison by Log-Expectation (MUSCLE) web tool was used to align protein sequences, enabling visualization of conserved residues using CLUSTALX. CLUSTALX outputs were aligned with a WEBLOGO frequency distribution plot.

### AlphaFold3 and Boltz-1 Protein Structure Modelling

Protein structure predictions of the HIF-2 dimer were generated to assess the structural impact of both a phosphomimetic mutation and phosphorylation of the HIF-2α protein compared to the wild type. ABCFold (Elliott *et al*., 2025) was used to run both AlphaFold3 (Abramson *et al*., 2024) and Boltz-1 (Wohlwend *et al*., 2024), as these software tools could predict structures with PTMs. ‘--mmseqs2’ and ‘--templates’ flags were added to use a the MMSeqs2 API for the Multiple Sequence Alignment (MSA) and the template searches (templates only used by AlphaFold3). The sequence used for the predictions was derived from the structure PDB:4ZP4 (Wu *et al*., 2015), chains A and B. The modifications applied to HIF-2α (PDB: 4ZP4:B) were as follows: phosphomimetic mutation S345 to S345D; phosphorylation S345 to S345SEP. The wild-type prediction remained unmodified.

## Statistical analysis

Data are expressed as mean± standard deviation. Comparisons among groups were made using ANOVA, with a Tukey correction where applicable. P values of <0.05 were considered significant. All data is based on a minimum of three independent biological replicates.

Venn diagrams generated using the BioVenn tool (Hulsen *et al*, 2008).

## Authors contributions

Conceptualization: AA, MB, VS, CEE, NSK, SR; Formal analysis: AA, LAD, KPF, SOO, GBE, NSK, SR; Investigation: AA, LAD, KPF, GB, LGE, NSK; Resources: JF, DJR, CEE, VS, NSK, SR; Writing original draft: AA, NSK, VS, CEE, SR; Writing reviewing and Editing: AA, LAD, KPF, SOO, GBE, GB, LGE, MB, DJR, JF, CEE, VS, NSK, SR; Visualisation: AA, NSK, GEB, GB, KPF, SOO, NSK, SR; Supervision: MB,CEE, NSK, VS, SR; Project administration: AA, SR; Funding acquisition: JF, DJR, CEE, NSK, VS, SR.

## Acknowledgements

We acknowledge the Liverpool Centre for Cell Imaging (CCI) for provision of imaging equipment, in particular the Epifluorescent microscope funded by the MRC grant number MR/K015931/1, the Cell Discovery 7 funded by the MRC grant number MR/X013502/1, OMERO and technical assistance. We thank the Imaging Center Essen (IMCES) at the Faculty of Medicine of the University of Duisburg-Essen, Germany for providing access to the Leica SP8 gSTED and FLIM. This work was funded by the Deutsche Forschungsgemeinschaft (DFG, German Research Foundation) – 234323630, INST 58219/26-1 FUGG. This work was funded by a Wellcome trust collaborator grant to SR (206293/Z/17/Z) and the University of Liverpool. AA was supported by a PhD studentship from the MRC. GB and GBE received a PhD studentship from the BBSRC. AA and NSK are supported by Worldwide Cancer Research [grant number 24-0353]. CEE was supported by a BBSRC grant (BB/R000182).

## Supplementary Information

Supplementary Figures, Tables and Datasets.

## Notes

### Competing Interest Statement

The authors have declared no competing interest.

