## Supplementary figures and tables for "Phosphorylation at S345 Converts HIF-2α from a Transcription Factor to an RNA binding protein"

Albanese et al, 2026:

**Phosphorylation at S345 Converts HIF-2α from a Transcription Factor to a Regulator of mRNA Fate**

**Supplementary information**

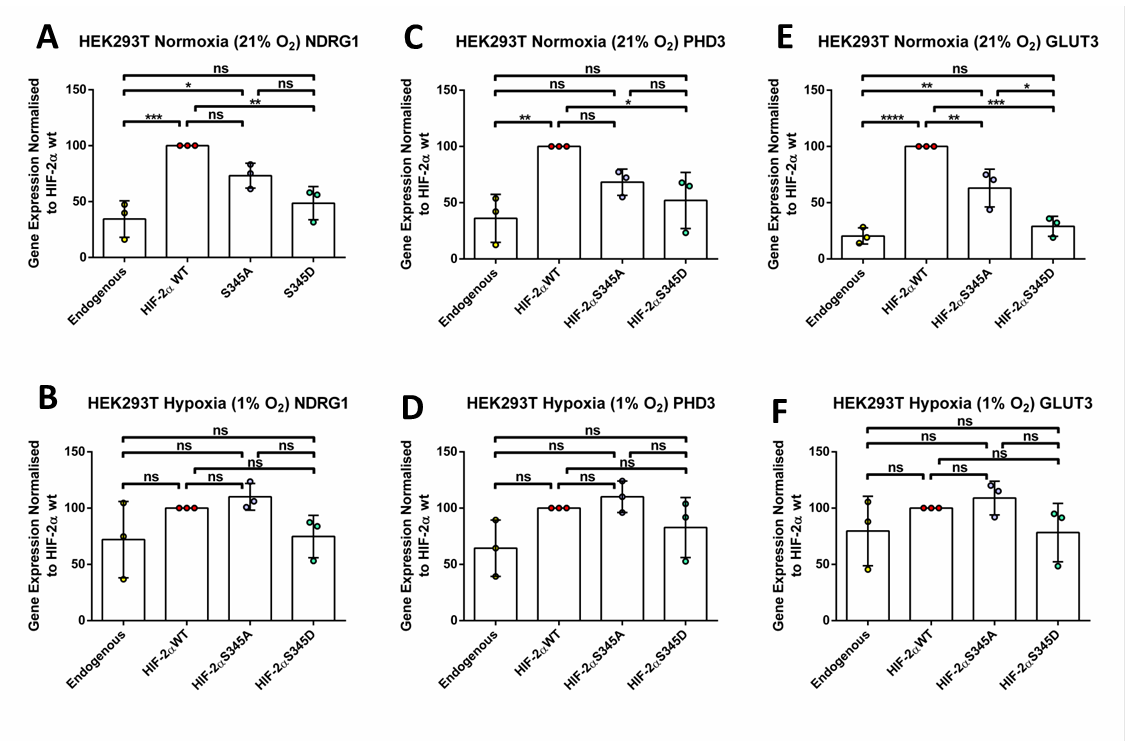

**Sup. Figure 1. HIF target induction in HEK293T cells determined at the gene level via RT-qPCR.** HEK293T cells were transiently transfected with either empty vector, HA-Clover HIF-2α, HA-Clover HIF-2α S345A or HA-Clover HIF-2α S345D for high level overexpression. (A and B) PHD3, (C and D) NDRG1 and (E and F) GLUT3, in normoxia and hypoxia (21% and 1% O2, respectively). All experiments were carried out as n = 3 independent experiments. Gene expression levels were normalised to HIF-2α wild-type. ±Standard deviation error bars are reported. One-way Anova Tukey post-hoc test was performed to determine significance (*P < 0.05; ns, not significant).

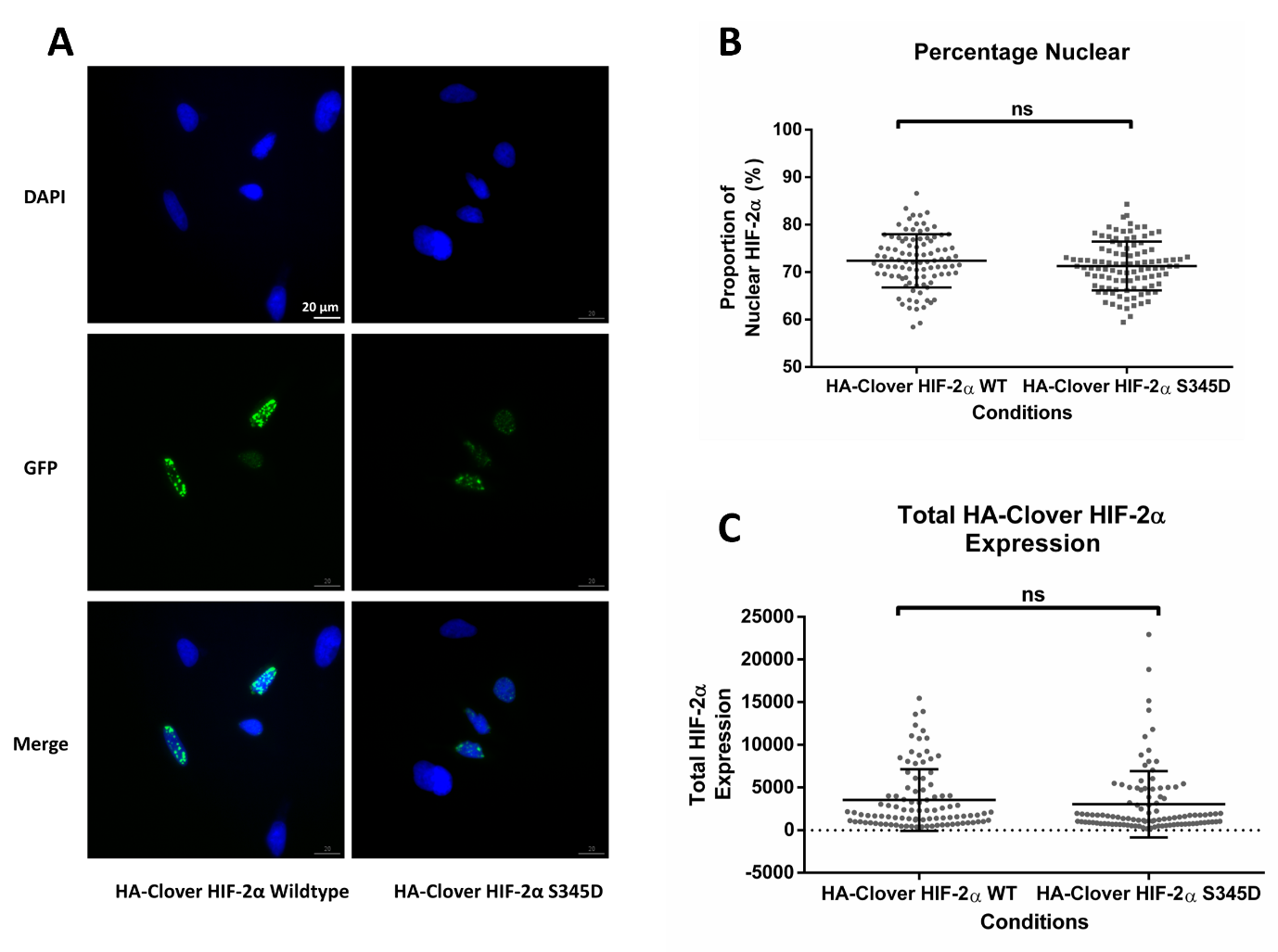

**Sup. Figure 2. Quantification of the intracellular accumulation of HIF-2α WT and S345D by 20x epifluorescent microscopy.** (A) Subcellular localisation of HA-Clover HIF-2α wild-type (WT) and S345D were assessed by fixed cell 20x epifluorescent microscopy. (B) The membrane permeable Hoechst stain was used to fluorescently label DNA to identify the nucleus and then imaged. Images of over 90 fixed HeLa cells per condition were obtained, from three replicates, to measure the proportion of nuclear HIF-2α. (C) Total exogenous HIF-2α levels for the WT and S345D mutant were determined by quantification of the GFP signal from the HA-Clover HIF-2α. 3 biological replicates were utilised, with each condition having a minimum of 30 cells imaged per biological replicate. A total of 97 and 104 cells were imaged for HIF2α WT and S345, respectively. A student T-test was used to determine statistical significance between conditions (P* <0.05, ns, not significant).

**Sup. Table 1. pTM scores from different alphaFold approaches**

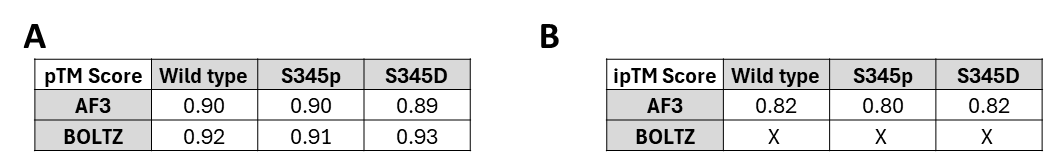

**Sup. Table 2. Unique hydrogen bonds of AlphaFold3 wild type, pS345, S345D and S345A HIF-2α within the HIF-2 heterodimers.** Chain A and B refer to HIF-1β and HIF-2α, respectively.

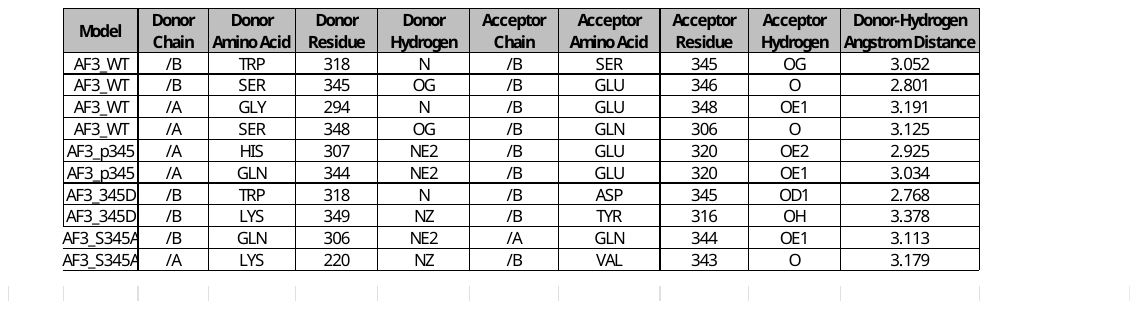

**Sup. Table 3. Overlap of proteins found in speckles (Alexander *et al*, 2025) and HIF-2α interactome.**

| Speckle proteins overlap with HIF-2α WT | Speckle proteins overlap with HIF-2α S345D |
| --- | --- |
| SF3B1 | TK1 |
| RBM39 | SNRNP40 |
| RPL24 | TNS3 |
| APRT | FLII |
|  | CAPZA2 |
|  | PCBP1 |
|  | THRAP3 |
|  | DDX17 |
|  | GIGYF2 |
|  | RAB34 |
|  | RPL13 |
|  | CDC37 |
|  | SKP1 |
|  | BCLAF1 |
|  | RFC3 |
|  | SIM1 |
|  | MYBBP1A |
|  | MYADM |
|  | SPTLC1 |
|  | ZNF572 |
